# Computing the Riemannian curvature of image patch and single-cell RNA sequencing data manifolds using extrinsic differential geometry

**DOI:** 10.1101/2021.01.08.425885

**Authors:** Duluxan Sritharan, Shu Wang, Sahand Hormoz

**Affiliations:** Harvard Graduate Program in Biophysics, Harvard University, Cambridge, MA, USA; Department of Data Sciences, Dana-Farber Cancer Institute, Boston, MA, USA; Laboratory of Systems Pharmacology, Harvard Medical School, Boston, MA, USA; Department of Systems Biology, Harvard Medical School, Boston, MA, USA; Broad Institute of MIT and Harvard, Cambridge, MA, USA

## Abstract

Most high-dimensional datasets are thought to be inherently low-dimensional, that is, datapoints are constrained to lie on a low-dimensional manifold embedded in a high-dimensional ambient space. Here we study the viability of two approaches from differential geometry to estimate the Riemannian curvature of these low-dimensional manifolds. The intrinsic approach relates curvature to the Laplace-Beltrami operator using the heat-trace expansion, and is agnostic to how a manifold is embedded in a high-dimensional space. The extrinsic approach relates the ambient coordinates of a manifold’s embedding to its curvature using the Second Fundamental Form and the Gauss-Codazzi equation. Keeping in mind practical constraints of real-world datasets, like small sample sizes and measurement noise, we found that estimating curvature is only feasible for even simple, low-dimensional toy manifolds, when the extrinsic approach is used. To test the applicability of the extrinsic approach to real-world data, we computed the curvature of a well-studied manifold of image patches, and recapitulated its topological classification as a Klein bottle. Lastly, we applied the approach to study single-cell transcriptomic sequencing (scRNAseq) datasets of blood, gastrulation, and brain cells, revealing for the first time the intrinsic curvature of scRNAseq manifolds.

## 1 Introduction

High-dimensional biological datasets have become prevalent in recent decades because of new technologies such as high-throughput scRNAseq [1, 2, 3], mass cytometry [4, 5] and multiplex imaging [6, 7]. Interpretation and visualization of such high-dimensional datasets have been challenging however, prompting the development of tools for non-linear projection of datapoints onto 2 or 3 dimensions [8]. These tools, such as IsoMAP [9], t-SNE [10] and UMAP [11], appeal to the ansatz that datapoints in a *high-dimensional ambient space* are constrained to lie on a *low-dimensional manifold*. Unfortunately, determining the geometry of a low-dimensional manifold from these visualizations is difficult, since many geometric properties are lost after projecting onto 2 or 3 dimensions. For example, the cartographic projections used in an atlas to flatten Earth’s curved surface tear apart continuous neighborhoods and non-uniformly stretch distances.

Fortunately, topology and differential geometry provide a wealth of concepts to characterize a manifold’s shape directly without confounding projections. In particular, *homology* [12, 13] categorizes a manifold according to the number of holes it contains, and the dimensionality of each hole (whereas for example, the hole in a hollow sphere does not survive projection onto a 2-dimensional plane). Similarly, *metrics* [14] and *geodesics* [9] determine shortest-distance paths between pairs of points on a manifold without any distortion from a projection (whereas for example, most atlases exaggerate distances at the poles). *Curvature* [15] is a local manifold property that quantifies the extent to which a manifold deviates from the tangent plane at each point *p*. Projecting a manifold onto a plane for visualization destroys this property by definition. Recent methods have emerged for estimating homology [16, 17], metrics [14] and geodesics [18] from noisy, sampled data, with accompanying statistical guarantees [18, 19, 20]. These methods have been applied to analyze images [21, 22] and biological datasets [23, 24]. However, estimating curvature has received less attention although it is fundamental to quantifying geometry.

Curvature arises from two sources. On the one hand, a manifold itself can be curved, resulting in *Riemannian* or *intrinsic curvature*. A sphere has intrinsic curvature because it cannot be flattened so that all geodesics on its surface correspond to straight lines on a Euclidean plane (see Figure 1A). On the other hand, the *embedding* of a manifold in an ambient space can give rise to *extrinsic curvature*, a property that is not inherent to the manifold itself. For example, a scroll has extrinsic curvature because it is formed by rolling a piece of parchment, but the parchment itself is not inherently curved (see Figure 1B). It is important to note that both types of curvature scale inversely with the global length scale (*L*) associated with a manifold. It is for this reason that a marble (*L* ≈ 1 cm) is visibly round, but the Earth (*L* ≈ 10, 000 km) is still mistaken by some to be flat. Since intrinsic curvature is an inherent property of a manifold, while extrinsic curvature is incidental to an embedding, we will restrict our attention to the former.

**Figure 1:**
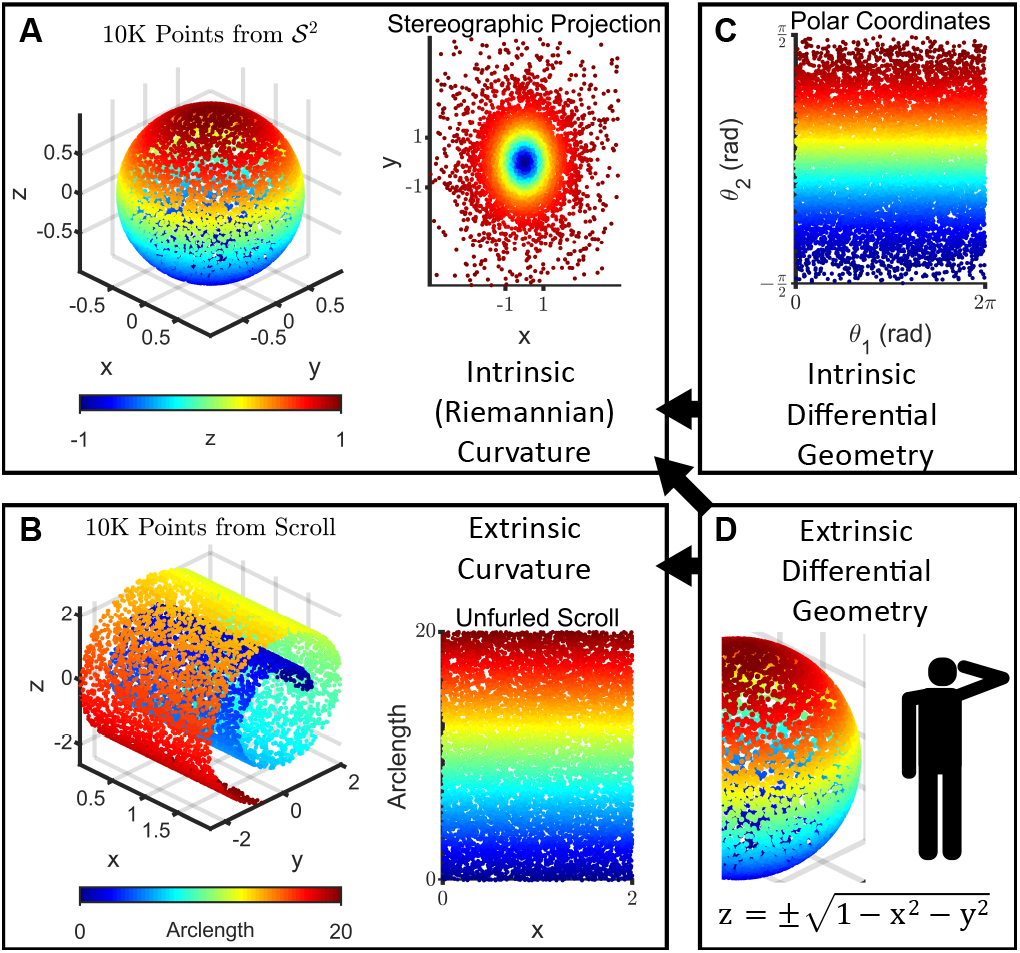
Riemannian curvature is an intrinsic property of a manifold while extrinsic curvature depends on the embedding. **(A)** (Left) *N* = 10^4^ points uniformly sampled from the 2-dimensional hollow unit sphere, 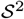, embedded in the 3-dimensional ambient space ℝ^3^, colored according to the z-coordinate. 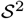 has Riemannian or intrinsic curvature because there is no projection onto 2-dimensional Euclidean space that preserves geodesic (shortest-path) distances. (Right) For example, a stereographic projection using the point *z* = (0, 0, 1) and the plane *z* = 0 introduces distortions since the geodesic distance between any pair of points in the lower hemisphere is (non-uniformly) larger than the Euclidean distance in this projection. **(B)** (Left) *N* = 10^4^ points uniformly sampled from a scroll, which is also a 2-dimensional manifold embedded in ℝ^3^. The scroll has extrinsic curvature because it curls away from the tangent plane at any point. (Right) However, it does not have intrinsic curvature, because it can be projected onto 2-dimensional Euclidean space in a way that preserves geodesic distances, by unfurling. **(C)** Intrinsic differential geometry treats manifolds as self-contained objects that can be described using only intrinsic coordinates, which do not depend on any embedding or ambient space. One possible set of intrinsic coordinates for 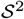 are polar coordinates, where *θ*_1_ and *θ*_2_ are the azimuthal and elevation angles respectively. While this representation superficially resembles the unfurled scroll in (B), distances in this plane are non-Euclidean. Any line segment along 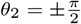 has zero length for example. **(D)** Extrinsic differential geometry defines manifolds in the coordinate system of the ambient space, which requires a privileged vantage point off the manifold itself. Both intrinsic and extrinsic differential geometry can be used to compute intrinsic curvature, whereas only extrinsic differential geometry can be used to compute extrinsic curvature (as indicated by the black arrows).

A precise description of intrinsic curvature is provided by the *Riemannian curvature tensor*, *R*_*lkij*_(*p*). For a given basis {*v*}, this tensor quantifies how much a vector initially pointing in direction *v*_*k*_ is displaced in direction *v*_*l*_ after parallel transport around an infinitesimal parallelogram defined by directions *v*_*i*_ and *v*_*j*_. The simplest intrinsic curvature descriptor is *scalar curvature*, *S*(*p*), which is formed by contracting *R*_*lkij*_(*p*) to a scalar quantity, as its name suggests. When *S*(*p*) is greater (less) than 0, the sum of the angles of a triangle formed by connecting three points near *p* by geodesics is greater (less) than *π*. Likewise, when *S*(*p*) is greater (less) than 0, a small ball centred at *p* has a smaller (larger) volume than a ball of the same radius in Euclidean space. We furnish toy examples in the main text to provide stronger intuition for this quantity.

In theory, intrinsic curvature can be equivalently computed using tools from either one of the two branches of differential geometry. *Intrinsic differential geometry* makes no recourse to an external vantage point off a manifold, just as the polygonal characters in Edwin Abbot’s classic Flatland [25] were confined to traversing in ℝ^2^, and found the notion of ℝ^3^ unfathomable. In this branch, a manifold is therefore represented in *intrinsic coordinates*, which are agnostic to any ambient space or embedding. A hollow sphere represented in polar coordinates and k-nearest neighbor (kNN) graph representations of a dataset, for instance, are in this spirit (see Figure 1C). Conversely, in *extrinsic differential geometry*, a manifold is treated as a surface embedded in an ambient space, and is represented in *ambient coordinates* (see Figure 1D). The surface of an organ is parameterized this way, for example, in a surgical robot suturing an incision.

In this work, we explore two approaches for estimating intrinsic curvature based on these twin views, keeping in mind practical limitations of real-world datasets, which may be comprised of a relatively small number of noisy measurements. The first approach uses the Laplace-Beltrami operator, which is well-studied in previous applications of differential geometry to data analysis [14, 26, 27, 28, 29], and is theoretically appealing as an intrinsic quantity that is embedding-invariant. However, we find that this approach cannot accurately estimate even average scalar curvature on the simplest of low-dimensional toy manifolds for small sample sizes, despite the history and ubiquity of the Laplace-Beltrami operator in geometric data analysis. Meanwhile, the second approach uses the Second Fundamental Form and the Gauss-Codazzi equation [15], identities that rely on information from the ambient space. We find that this extrinsic approach is not only more robust to small sample sizes and noise, but permits computation of the full Riemannian curvature tensor, though we focus on the scalar curvature for simplicity. Using these insights, we developed a software package to compute the scalar curvature (and associated uncertainty) at each sampled point on a manifold, and applied this tool to investigate the curvature of image and scRNAseq datasets.

## 2 Results

### 2.1 Estimators of the Laplace-Beltrami Operator Yield Inaccurate Scalar Curvatures

Intrinsic differential geometry treats a *d*-dimensional manifold, *M*, as a self-contained object and is agnostic to how *M* may be represented in ambient coordinates due to any particular embedding (see Figure 1C). Conceptually, this is accomplished by only considering *M* as a collection of local, overlapping neighborhoods. The geometry of these neighborhoods is encoded using tools such as the Laplace-Beltrami operator, Δ_*M*_, which captures diffusion dynamics across neighborhoods. For most practical applications, we do not have direct access to *M* but instead to a finite number (*N*) of points sampled from *M*. For these cases, estimators of Δ_*M*_ are used instead. These estimators are well-studied [14, 26, 27, 28, 29], and the convergence rates of some have been characterized [30].

The scalar curvature averaged across *M*, has a well-known connection to Δ_*M*_ via the heat-trace expansion [27, 31], which relates the eigenvalues, *λ*_*k*_, of Δ_*M*_ to the geometry of *M* :

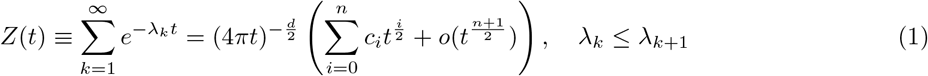

The first few coefficients, *c*_*i*_, are given by [27]:

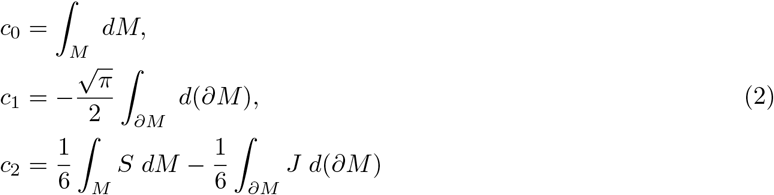

where *∂M* is the boundary of the manifold and *J* is the mean curvature on *∂M*. Recall that *S* is the point-wise scalar curvature. By inspection, *c*_0_ is the volume, *c*_1_ is proportional to the area, and *c*_2_ is directly related to the average scalar curvature.

We reasoned that if the average scalar curvature cannot be accurately computed for a manifold with constant scalar curvature using these relations, then computing the point-wise scalar curvature for more complex manifolds is intractable. To investigate this, we considered the 2-dimensional hollow unit sphere, 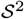, for which the true scalar curvature is *S*(*p*) = 2 ∀*p* ∈ *M*, and uniformly sampled *N* = 10^4^ points to mirror the typical size of current scRNAseq datasets (see Figure 1A; Methods Section 4.4.1.1).

Since common estimators of Δ_*M*_ only yield as many eigenvalues as datapoints (*N*), we cannot compute the infinite set of eigenvalues needed in Equation 1. Therefore, we introduced a truncated series with *m* eigenvalues, *z*_*m*_(*x*), where we have substituted 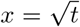 and divided through by the prefactor in the RHS of Equation 1 to isolate for *c*_*i*_, following the approach in [27]:

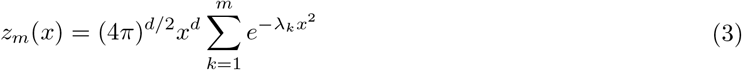

The scalar curvature can then be approximated by fitting the truncated series, *z*_*m*_(*x*), to a second-order polynomial, *p*_2_(*x*), over intervals of small *x*:

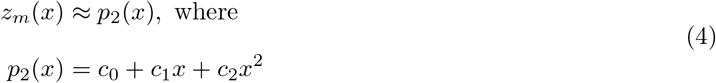

We estimated Δ_*M*_ using the *N* sampled points (see Methods Section 4.2.5), substituted the eigenvalues of the estimate into Equation 3, and numerically fit *z*_*m*_(*x*) to *p*_2_(*x*) (see Figure S1A-G; Methods Section 4.2.1). We obtained the scalar curvature by inspecting the resulting *c*_2_ coefficient, and compared the result to the true value of 2. We found that the scalar curvature was always over-estimated (*S* > 3) regardless of *m*, the number of eigenvalues used in the truncated series (see Methods Section 4.2.3), or the choice of estimator for Δ_*M*_ (see Methods Section 4.2.5). We identified the poor convergence of the estimated eigenvalues of Δ_*M*_ as the source of error (see Methods Section 4.2.4) and found that at least *N* ≈ 10^7^ points are required to reduce the error to ±0.5, so that *S* ≈ 2.5 (see Figure S1H).

Therefore, despite the prevalence of the Laplace-Beltrami operator in geometric data analysis, our example shows that an intrinsic approach relying on the operator is not practical for computing scalar curvatures. Even for noise-free datapoints uniformly sampled from 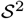, the sample size needed to compute average scalar curvature accurate to ±0.5 is several orders of magnitude greater than what is typically feasible in current scRNAseq experiments. Noise and non-uniform sampling would confound the issue further. Most importantly, we would eventually like to compute local values of *S*(*p*) ∀*p* ∈ *M*, but this approach failed to correctly recover even average scalar curvature, which one might have expected to be feasible. To find an alternative approach, we next considered tools from extrinsic differential geometry.

### 2.2 Curvature Can Be Computed Accurately Using the Second Fundamental Form

In extrinsic differential geometry, a manifold is described in the coordinates of the ambient space in which it is embedded, usually ℝ^*n*^ (see Figure 1D). Since the shape of the sphere in Figure 1A is visually unambiguous to the eye (thanks to its extrinsic view from a vantage point off the manifold), we reasoned that an extrinsic approach would be more fruitful.

A *d*-dimensional manifold, *M*, embedded in ℝ^*n*^ can be described at each point *p* in terms of a *d*-dimensional tangent space, *T*_*M*_ (*p*), and an (*n* − *d*)-dimensional normal space, *N*_*M*_ (*p*), as shown in Figure 2A. Given orthonormal bases for *T*_*M*_ (*p*) and *N*_*M*_ (*p*), points in the neighborhood of *p* can be expressed as *Y* = [*t*_1_,…, *t*_*d*_, *n*_1_,…, *n*_*n*−*d*_] where *t*_*i*_ is *Y* ’s coordinate along the *i*^th^ basis vector of *T*_*M*_ (*p*) and *n*_*k*_ is *Y* ’s coordinate along the *k*^th^ basis vector of *N*_*M*_ (*p*). The *n*_*k*_s can then be locally approximated as functions of the *t*_*i*_s i.e. *n*_*k*_ ≈ *f*_*k*_(*t*_1_,…, *t*_*d*_) as shown in Figure 2B.

**Figure 2:**
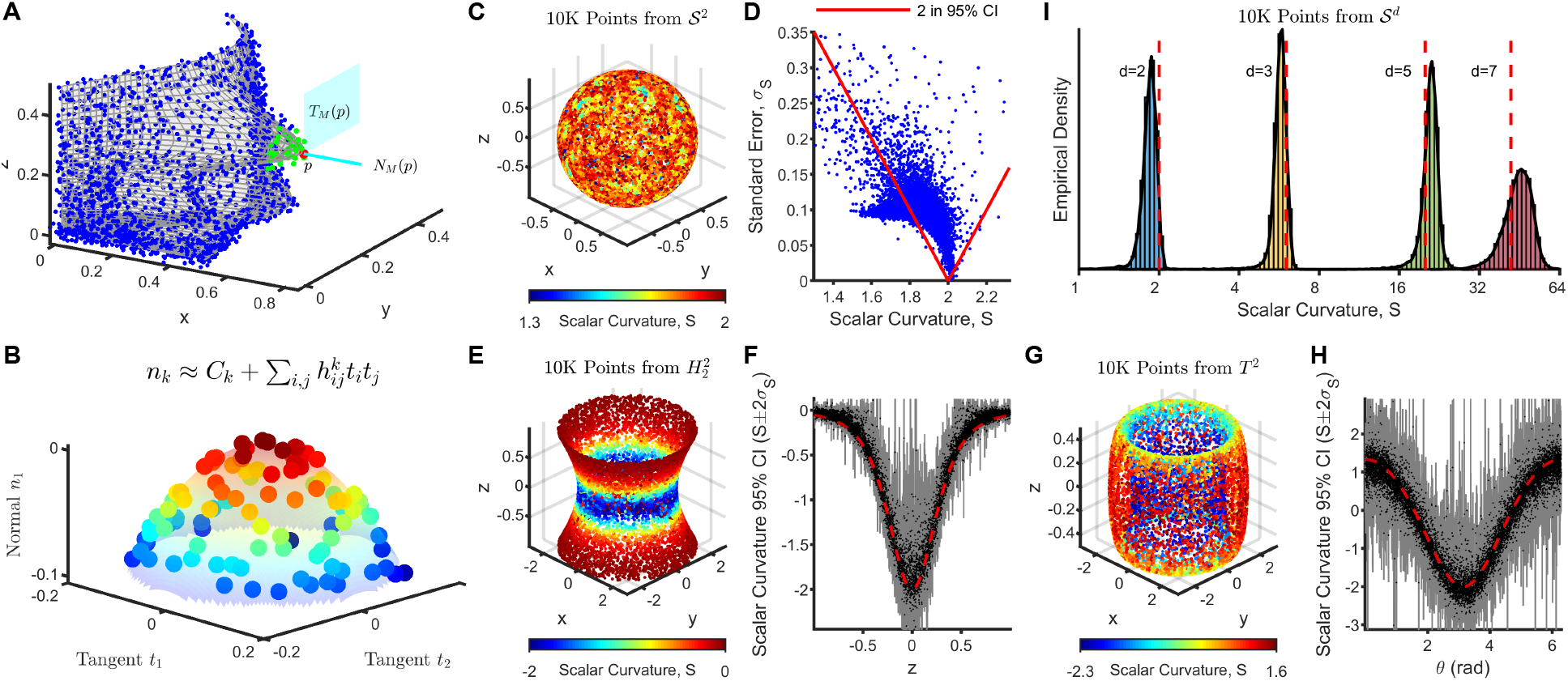
Scalar curvature is accurately estimated using the Second Fundamental Form and the Gauss-Codazzi equation. **(A)** A hypothetical manifold (shown in grey) from which datapoints are sampled (shown as colored dots). The manifold at any given point *p* (shown in red) can be decomposed into a tangent space *T*_*M*_ (*p*) (the cyan plane) and a normal space *N*_*M*_ (*p*) (the cyan line). Points in the neighborhood around *p* (shown in green) can be expressed in terms of orthonormal bases for *T*_*M*_ (*p*) and *N*_*M*_ (*p*) (see (B) below). **(B)** The set of points in the neighborhood of *p* (shown as green dots in (A)) are represented here in local tangent (*t*_1_*, t*_2_) and normal (*n*_1_) coordinates, corresponding to orthonormal bases for *T*_*M*_ (*p*) and *N*_*M*_ (*p*) respectively. Coloring corresponds to magnitude in the normal direction. The normal coordinates (*n*_1_) can be locally approximated as a quadratic function (the translucent surface) of the tangent coordinates (*t*_1_*, t*_2_), according to the Second Fundamental Form, 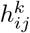. **(C)** Scalar curvatures computed using the extrinsic approach for *N* = 10^4^ points uniformly sampled from the 2-dimensional hollow unit sphere, 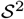. The true value is 2 at all points on the manifold. See Methods Section 4.4.1.1. **(D)** Scalar curvatures (*S*) computed in (C) are plotted against their associated standard errors (*σ*_*S*_). Points enclosed by the red lines have a 95% confidence interval (CI), computed as *S* ± 2*σ*_*S*_, containing the true value of 2. **(E)** As in (C) but for *N* = 10^4^ points uniformly sampled from a one-sheet hyperboloid, 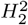, which is also a 2-dimensional manifold. Due to the radial symmetry of the manifold, scalar curvature only varies only along the *z*-direction. See Methods Section 4.4.1.2. **(F)** Scalar curvatures (black) computed in (E) with their associated 95% CIs (shown in grey) plotted as a function of the z-coordinates of the datapoints. The true value is shown as a dashed red line. **(G)** As in (C) but for *N* = 10^4^ points uniformly sampled from a 2-dimensional ring torus, *T* ^2^. *T* ^2^ is constructed by revolving a circle parameterized by *θ*, oriented perpendicular to the *xy*-plane, through an angle *ϕ* around the *z*-axis. The scalar curvature only depends on the value of *θ*. See Methods Section 4.4.1.3. **(H)** Scalar curvatures computed in (G) with their associated 95% CIs plotted as a function of the *θ* values of the datapoints. Colors as in (F). **(I)** Distribution of computed scalar curvatures for *N* = 10^4^ points uniformly sampled from the *d*-dimensional unit hypersphere, 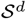, for *d* = 2, 3, 5, 7. As with 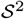, these manifolds are isotropic and have constant scalar curvature. The true values are shown as dashed red lines. See Methods Section 4.4.1.1.

The Riemannian curvature of *M* is related to the quadratic terms in the Taylor expansion of each *f*_*k*_ with respect to the *t*_*i*_s. Specifically, the *Second Fundamental Form* of *M*, 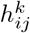, gives the second-order coefficient relating each *f*_*k*_ to the quadratic term *t*_*i*_*t*_*j*_ [32]:

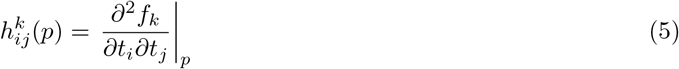

The Riemannian curvature tensor is related to the Second Fundamental Form according to the Gauss-Codazzi equation [15]:

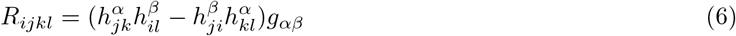

where *g*_*αβ*_ is the metric of the ambient space, which we take to be the usual Euclidean metric *δ*_*α,β*_ going forward. The scalar curvature can be obtained by contracting the Riemannian curvature tensor:

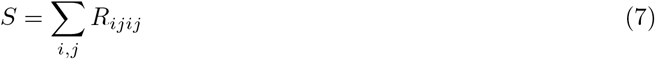

This suggests a conceptually simple procedure to estimate the scalar curvature of a data manifold at each point *p*: (i) estimate *T*_*M*_ (*p*) and *N*_*M*_ (*p*), (ii) determine 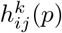 in local coordinates, (iii) compute *S* using Equations 6 and 7. We developed a computational tool that provides an implementation of this procedure. Briefly, given a set of datapoints {*X*} ∈ ℝ^*n*^ and manifold dimension *d*, a neighborhood around each point *p* is selected to be the *n*-dimensional ball centred on *p* of radius *r* encompassing *N*_*p*_(*r*) points (see Methods Section 4.3.2). For each point *p*, Principal Component Analysis (PCA) [33] is performed on the *N*_*p*_(*r*) points in its neighborhood, and the first *d* (last *n* − *d*) Principal Components (PCs) accounting for the most (least) variance are taken as an orthonormal basis for *T*_*M*_ (*p*) (*N*_*M*_ (*p*)). The normal coordinates, *n*_*k*_, of the *N*_*p*_(*r*) points in each neighborhood are fit by regression to a quadratic model in terms of the tangent coordinates, *t*_*i*_, to obtain 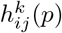 with associated uncertainties (see Figure 2B; Methods Section 4.3.1).

The choice of *r*(*p*) is an important one since it sets the length scale at which curvature is computed for point *p* (see Methods Section 4.3.5). Our tool allows interrogation of curvature at any length scale of interest by allowing the user to manually set *r*(*p*), a feature we use to inspect real-world datasets later in the paper. However, since the local geometry of the manifold may be non-trivial and unknown *a priori*, we also provide the ability to set *r*(*p*) according to statistical rather than geometric principles. Specifically, our tool algorithmically chooses *r* at each *p* so that the uncertainty in 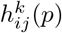 from regression is less than a user-specified global parameter, *σ*_*h*_ (see Methods Section 4.3.2). Since a larger number of points reduces the uncertainty in regression, a smaller *σ*_*h*_ requires a larger *r*(*p*) ∀*p* ∈ *M*. This strategy of setting *σ*_*h*_ therefore allows neighborhood sizes to dynamically vary over the manifold based on the local density of the data, which means the algorithm can gracefully handle non-uniform sampling of the manifold. The choice of *σ*_*h*_ will depend on the global length scale, *L*, of the datapoints (see Methods Section 4.3.5), the average density of sampled points, and of course, the desired uncertainty in the estimates of 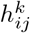. These uncertainties are in turn used to compute a standard error, *σ*_*S*_, accompanying the scalar curvature estimate at each point, using standard error propagation formulas (see Methods Section 4.3.4). We specify *σ*_*h*_ instead of *σ*_*S*_ as the global parameter for choosing neighborhood sizes, since the latter depends non-linearly on the values of 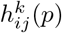, which makes determining *r*(*p*) more difficult.

Our algorithm also computes a goodness-of-fit (GOF) p-value at each *p* by comparing residuals from regression against a normal distribution to quantify how well the normal coordinates are fit by a quadratic function (see Methods Section 4.3.3). We tested this p-value at significance level *α* = 0.05, declaring fits to be poor when the residuals are significantly non-Gaussian. The p-value can be disregarded if the neighborhood size is manually specified to be larger than a length scale for which a quadratic fit is appropriate. However, when *σ*_*h*_ is specified instead, a uniform distribution of these p-values over *M* indicates that the desired uncertainty results in neighborhoods that are well-approximated using quadratic regression. We adopted this heuristic when choosing *σ*_*h*_ for the datasets studied in this paper (see Methods Section 4.4.3, 4.5.7 and 4.6.4). The software is available at https://gitlab.com/hormozlab/ManifoldCurvature.

We first applied our algorithm to compute scalar curvatures for the same *N* = 10^4^ points uniformly sampled from 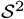 for which the intrinsic approach failed (see Figure 2C; Methods Section 4.4.1.1). The algorithm yielded scalar curvature estimates at each point with mean error −0.17 (computed by averaging the difference between the point-wise scalar curvature estimates and the ground truth value of 2 across all points) using neighborhoods that only contained *N*_*p*_(*r*) ≈ 10^2^ points. This is already superior to the intrinsic approach, which failed to compute even average scalar accurate to ±1 for the same sample size. The non-zero value of the mean error indicates that our estimator is biased. The values of 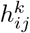 are not biased because they are obtained using regression. Even so, the components of the Riemannian curvature tensor, *R*_*ijkl*_, may still be biased because they are non-linear functions of 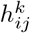. Note that for 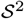, this bias is the same across all datapoints (because of the isotropic nature of the manifold) and therefore results in a systematic under-estimation of scalar curvature (see Figure 2C; Methods Sections 4.3.4). We also computed 95% confidence intervals (CI) for our estimates as *S* ± 2*σ*_*S*_, and despite the mean error, 73% of points still reported a 95%CI containing the true value of 2 (see Figure 2D).

We next tested our algorithm on a 2-dimensional manifold with negative scalar curvature, by uniformly sampling *N* = 10^4^ points from the one-sheet hyperboloid, 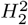 (see Figure 2E; Methods Section 4.4.1.2). Here, 71% of points reported a 95% CI containing the true scalar curvature (see Figure 2F). Lastly, we considered the 2-dimensional ring torus, *T* ^2^ (see Figure 2G; Methods Section 4.4.1.3). As a manifold with regions of positive, zero, and negative scalar curvature, *T* ^2^ is a useful toy model for understanding more complex 2-dimensional manifolds and gaining intuition for higher-dimensional manifolds. In 2 dimensions, regions of a manifold with positive scalar curvature (*θ* = 0, 2*π* in Figure 2H) are dome-shaped, regions with zero scalar curvature (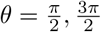 in Figure 2H) are planar, and regions with negative scalar curvature (*θ* = *π* in Figure 2H) are saddle-shaped. We applied our tool to *N* = 10^4^ points uniformly sampled from *T*^2^ and found that 88% of points reported a 95% CI containing the true scalar curvature (see Figure 2H).

To test the applicability of our algorithm to higher-dimensional manifolds, we uniformly sampled *N* = 10^4^ points from unit hyperspheres, 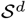, and found that 90%, 84% and 78% of points reported a 95% CI containing the true scalar curvature for *d* = 3, 5 and 7 respectively (see Figure 2I; Methods Section 4.4.1.1). The number of terms, 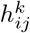, in the Second Fundamental Form grows as *d*^2^. For larger *d*, a greater number of datapoints and hence larger neighborhoods are needed for regression, but these are no longer well-approximated by quadratic fits according to our GOF measure. More generally, higher-dimensional manifolds require a higher density of data to estimate scalar curvatures accurately.

We additionally characterized how our algorithm performed when datapoints were non-uniformly sampled (see Figure S2A; Methods Section 4.4.2.1) or convoluted by observational noise (see Figure S2B; Methods Section 4.4.2.2), when the dimension of the ambient space was large (see Figure S2C; Methods Section 4.4.2.3), and when the specified manifold dimension differed from the ground truth (see Figure S2D; Methods Section 4.4.2.4). We found that the algorithm is robust to non-uniform sampling, large ambient dimension and small observational noise, and provides signatures indicating when the manifold dimension may be misspecified. However, when the noise scale is large, the resulting manifold is no longer trivially related to the noise-free manifold, consistent with existing literature [34, 35, 36, 37], so that scalar curvature cannot be accurately computed. Lastly, we note that since the full Riemannian curvature tensor is computed as an intermediate step in our algorithm, more intricate geometric features in the data can also be analyzed using our tool, though we defer such investigation to future studies.

Taken together, these examples demonstrate the utility of the algorithm in recovering curvature with specified uncertainties for manifolds with positive and/or negative scalar curvature. Next, we tested our algorithm on real-world data.

### 2.3 Curvature of Image Patch Manifold is Consistent with a Noisy Klein Bottle

Pixel intensity values in images of natural scenes are not independently or uniformly distributed. Understanding the statistics of such images is important for designing compression algorithms [38] and for addressing challenges in the field of computer vision such as segmentation [39]. Lee et al. discovered that 3×3-pixel patches extracted from greyscale images of natural scenes, whose pixels have high-contrast (i.e. the differences between the intensity values of adjacent pixels in a patch are large), are not uniformly distributed in ℝ^9^, but are instead concentrated on a low-dimensional manifold [40]. This is because high-contrast regions in a natural scene usually correspond to the edges of objects in the scene. High-contrast image patches consequently tend to be comprised of gradients and not simply random speckle. Subsequent work using topological data analysis revealed that after appropriate normalization (which takes image patches from ℝ^9^ to 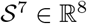, so that the global length scale is *L* = 1; see Methods Section 4.5.2), dense regions of high-contrast image patches have the same homology as a 2-dimensional manifold called a Klein bottle [21].

A Klein bottle, *K*^2^, is a canonical manifold typically introduced in the context of orientability, where it is often visualized in ℝ^3^ (as shown in Figure 3A) to highlight that it is non-orientable. From a topological perspective, *K*^2^ is a manifold parameterized by *θ, ϕ* ∈ [0, 2*π*] as shown in Figure 3B in which vertical edges are defined to be *θ* = 0 and *θ* = *π*, and horizontal edges are defined to be *ϕ* = 0 and *ϕ* = 2*π*. To make a closed surface, the vertical (horizontal) edges are glued together according to the red (blue) arrows in Figure 3B. *K*^2^ is therefore 2*π*-periodic in *ϕ*, since a point corresponding to *θ* on the bottom horizontal edge (*ϕ* = 0) is the same as the point corresponding to *θ* on the top horizontal edge (*ϕ* = 2*π*). Similarly, a point corresponding to *ϕ* on the left vertical edge (*θ* = 0) is the same as the point corresponding to 2*π* − *ϕ* on the right vertical edges (*θ* = *π*). In short, points on *K*^2^ obey the similarity relation (*θ, ϕ*) ∼ (*θ* + *π,* 2*π* − *ϕ*). *K*^2^ captures the dominant features in high-contrast image patches because *θ* can be treated as a parameter controlling rotation and *ϕ* as a parameter controlling the relative contribution of linear vs. quadratic gradients (see Figure 3B).

**Figure 3:**
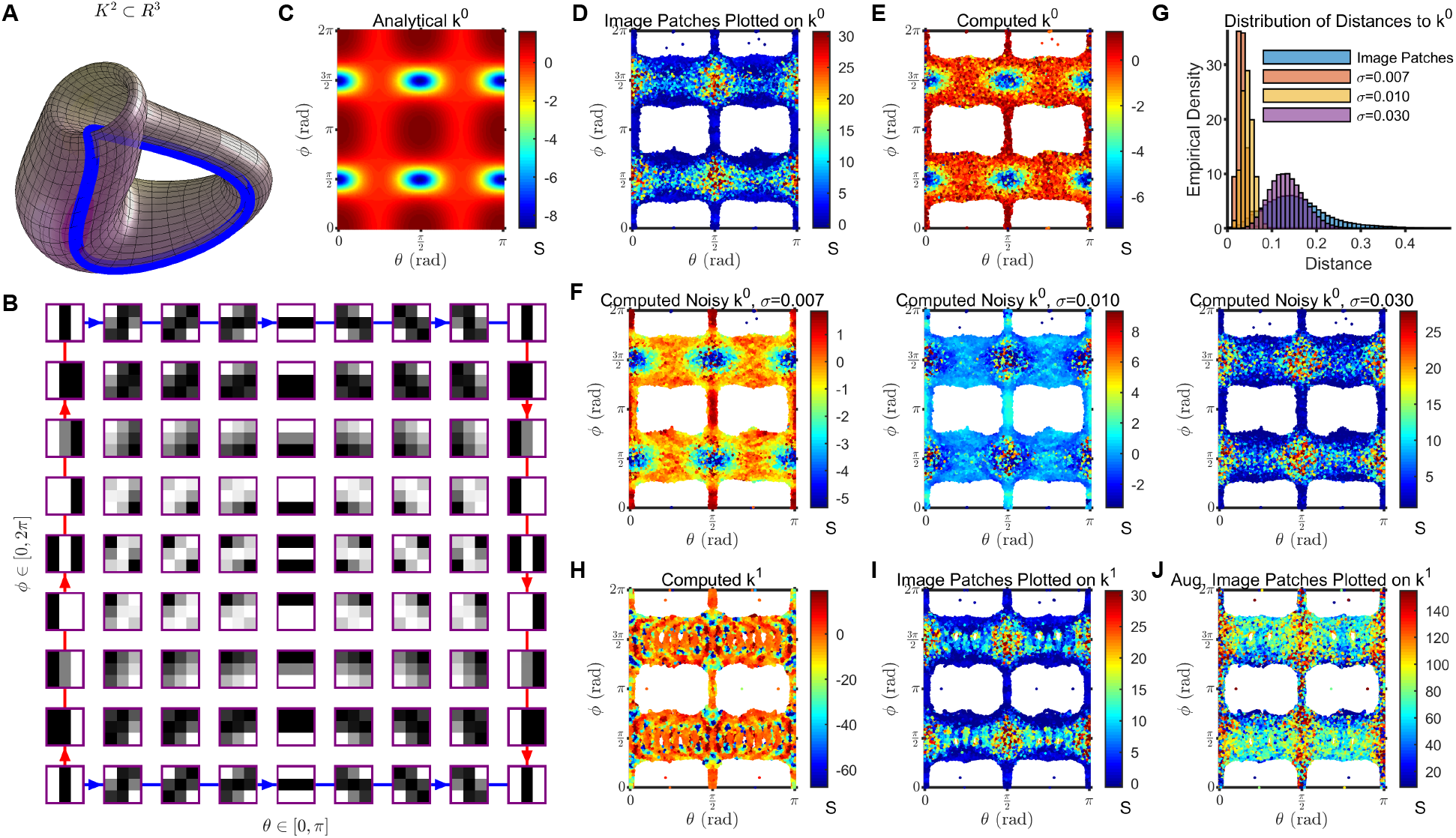
Scalar curvature computed for image patches is consistent with that of a Klein bottle with added isotropic Gaussian noise. **(A)** The Klein bottle, *K*^2^, is a 2-dimensional manifold shown here in ℝ^3^. **(B)** *k*^0^ is an analytical embedding given by Carlsson et al. in [21] relating parameter values *θ, ϕ* [0, 2*π*] to 3×3-pixel patches of greyscale images (see Equation 31 in Methods Section 4.5.3). *θ* controls the rotation of stripes in the image patches and *ϕ* determines the relative contribution of linear vs. quadratic gradients. Importantly, as shown in the figure, this embedding has boundary conditions consistent with the topology of a Klein bottle (depicted by the blue/red arrows). In particular, the embedding produces image patches that obey the similarity relation (*θ, ϕ*) ~ (*θ* + *π,* 2*π ϕ*). Adapted from Figure 6 of [21]. **(C)** The analytical scalar curvature of *k*^0^ (derived as described in Methods Section 4.1). **(D)** Scalar curvatures computed for *N* ≈ 4.2 10^5^ high-contrast 3×3-pixel patches sampled from the greyscale images in the *van Hateren* dataset [41] are plotted here as a function of (*θ*_0_*, ϕ*_0_), the parameter values of the closest point on *k*^0^ associated with each image patch (see Methods Section 4.5.4). **(E)** Scalar curvatures computed for the set of *N* ≈ 4.2 10^5^ closest points on *k*^0^ associated with the image patches. Note the close correspondence with Figure 3C, indicating that our algorithm correctly recapitulates the analytical scalar curvature. **(F)** As in (E), but after adding isotropic Gaussian noise in ℝ^9^ to the set of closest points on *k*^0^ (see Methods Section 4.5.6). Left to right corresponds to increasing levels of noise, *σ* = 0.007, 0.01, 0.03. **(G)** The distribution of Euclidean distances in ℝ^8^ between each image patch and its closest point on *k*^0^ is shown in blue. The distribution of distances to *k*^0^ after adding Gaussian noise to these closest points on *k*^0^ is also shown. **(H)** *k*^1^ is the analytical embedding from *θ, ϕ* [0, 2*π*] to ℝ^9^ that minimizes the sum of Euclidean distances from the image patches to the closest point on the embedding (see Methods Section 4.5.5). Each of the *N* ≈ 4.2 10^5^ image patches was associated to its closest point on *k*^1^, given by parameter values (*θ*_1_*, ϕ*_1_) (see Methods Section 4.5.4). Scalar curvatures computed on this set of *N* ≈ 4.2 × 10^5^ points on *k*^1^ are shown. **(I)** The same scalar curvatures computed for the image patches and visualized on (*θ*_0_*, ϕ*_0_) coordinates in (D), are shown here plotted on (*θ*_1_*, ϕ*_1_) coordinates. **(J)** Scalar curvatures computed for a densely sampled manifold comprised of the full set of *N* ≈ 1.3×10^8^ high-contrast 3×3-pixel image patches in the *van Hateren* image dataset (see Methods Section 4.5.2), visualized on (*θ*_1_*, ϕ*_1_) coordinates.

An embedding of *K*^2^ into ℝ^9^ with an analytical form, *k*^0^, was proposed by Carlsson et al. in [21] to model image patches (see Equation 31 in Methods Section 4.5.3). This embedding takes points from (*θ, ϕ*) into image patches in ℝ^9^ as shown in Figure 3B. For example, 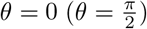 corresponds to patches with vertical (horizontal) stripes and 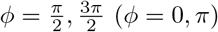 corresponds to patches with linear (quadratic) gradients. As *θ* increases, stripes in the image patches are rotated clockwise. As *ϕ* increases, image patches oscillate between having quadratic and linear gradients. Importantly, the image patches constructed by this embedding obey the same similarity relation (*θ, ϕ*) ∼ (*θ* +*π,* 2*π* −*ϕ*) topologically required of a Klein bottle. Whereas Carlsson et al. studied the global topology of image patches using this embedding, here we study their local geometry instead.

First, we analytically calculated the scalar curvature of *k*^0^ as a function of (*θ, ϕ*) as shown in Figure 3C (see Methods Section 4.1). Next, we used our algorithm to compute the scalar curvature on a data manifold of *N* ≈ 4.2×10^5^ high-contrast 3×3-pixel image patches randomly sampled from the same *van Hateren* dataset used to propose *k*^0^ (see Methods Section 4.5.2). We picked *σ*_*h*_ so that the distribution of GOF p-values was flat, and fixed this value for all subsequent simulations (see Methods Section 4.5.7). To visualize the results, we associated each image patch to its closest point on *k*^0^ (see Methods Section 4.5.4), and plotted the scalar curvatures on the resulting (*θ*_0_*, ϕ*_0_) coordinates (see Figure 3D). Most image patches map to 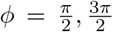 or 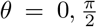 because linear gradients (of any orientation) and quadratic gradients that are vertically or horizontally oriented are the dominant features in the data as previously reported [21, 40].

The scalar curvatures computed for the image patches did not match the analytical scalar curvature of *k*^0^ (cf. Figures 3C and 3D). To identify the cause of this discrepancy, we first validated our algorithm by computing scalar curvatures on the set of *N* ≈ 4.2 × 10^5^ (*θ*_0_*, ϕ*_0_) points on *k*^0^ associated with the image patches (see Figure 3E); we found close agreement with the analytical calculation (75% of points reported a 95% CI containing the true scalar curvature). Next, observing that the neighborhood sizes used for computing the scalar curvature of image patches were larger than those used for computing the scalar curvature of the associated (*θ*_0_*, ϕ*_0_) points (cf. Figures S3A and S3B), we recomputed the scalar curvatures of these (*θ*_0_*, ϕ*_0_) points, but now with the same neighborhood sizes used for the image patches. The results agreed with the analytical calculation, but still did not match the scalar curvatures computed for the image patches (see Figure S3C).

Having ruled out these two possibilities, we hypothesized that the discrepancy was caused by fluctuations in the positions of the image patches with respect to the (*θ*_0_*, ϕ*_0_) points on the *k*^0^ manifold (real image patches are noisy and the Klein bottle embedding is only an idealization). We found that adding isotropic Gaussian noise of increasing magnitude in ℝ^9^ to the set of (*θ*_0_*, ϕ*_0_) points on *k*^0^ indeed resulted in scalar curvatures that resemble the data (see Figure 3F; Methods Section 4.5.6). The best agreement between the scalar curvatures of the image patches and the noisy (*θ*_0_*, ϕ*_0_) points was achieved when the magnitude of noise was *σ* = 0.03. Notably, in this case, the median Euclidean distance of the noisy (*θ*_0_*, ϕ*_0_) points to *k*^0^ was 0.132, which is comparable to 0.148, the median Euclidean distance of the image patches to *k*^0^ (see Figure 3G). Furthermore, the neighborhood sizes chosen by our algorithm when *σ* = 0.03 (see Figure S3A) matched those chosen for the image patches (see Figure S3B).

To find an embedding of the Klein bottle that might better explain the scalar curvature of the image patches without needing to add noise, we incorporated higher-order terms to *k*^0^ (see Methods Section 4.5.3). The coefficients for the higher-order terms were determined by fitting the data, resulting in a new embedding, which we refer to as *k*^1^ (see Methods Section 4.5.5). The median Euclidean distance of the image patches to *k*^1^ was 0.115 versus 0.148 to *k*^0^. As was done for *k*^0^, we associated each image patch to its closest point (*θ*_1_*, ϕ*_1_) on *k*^1^, and used our algorithm to compute the scalar curvature of these (*θ*_1_*, ϕ*_1_) points (see Figure 3H). Despite the reduction in the median Euclidean distance of images patches to the embedding, the scalar curvature of *k*^1^ was even less similar to that of the image patches (visualized in Figure 3I on these new (*θ*_1_*, ϕ*_1_) coordinates for *k*^1^) than was the scalar curvature of *k*^0^; the range of scalar curvature values for *k*^1^ was much larger than for either the image patches or *k*^0^, and the scalar curvature fluctuates on smaller length scales.

Lastly, we reasoned that there might be fine-scale scalar curvature fluctuations in the image patches that are masked by the larger neighborhood sizes used to compute scalar curvature for the image patches (see Figure S3B) relative to *k*^1^ (see Figure S3D). To decrease the neighborhood sizes chosen by the algorithm for the same *σ*_*h*_, we augmented the image patch dataset using the full set of *N* ≈ 1.3 × 10^8^ datapoints from the *van Hateren* dataset (see Methods Section 4.5.2). This resulted in neighborhood sizes comparable to those determined for *k*^1^ (cf. Figures S3D and S3E), but failed to recapitulate the fine-scale scalar curvature fluctuations observed in *k*^1^ (see Figure 3J). As a sanity check, we confirmed that the scalar curvature of the augmented image patch dataset matched that of the original image patch dataset, when computed using the same neighborhood sizes as the latter (see Figure S3F). Therefore, including higher-order terms in the embedding does not yield scalar curvatures that better agree with the data. Taken together, our analysis of curvature suggests that the image patch dataset can be best modelled by adding noise to the simplest embedding, *k*^0^.

Having applied our algorithm on real-world manifold-valued data that is well-modelled by an analytical embedding, we next turned our attention to scRNAseq datasets, which are generally regarded as low-dimensional manifolds and have no known analytical form.

### 2.4 scRNAseq Datasets have Non-Trivial Intrinsic Curvature

In scRNAseq datasets, each datapoint corresponds to a cell, and each coordinate to the abundance of a different gene. Here we consider the data manifold after basic preprocessing and linear dimensionality reduction using PCA (see Methods Section 4.6.1). Since many common analyses in the field such as clustering, visualization, and inference of cell differentiation trajectories are performed in this reduced space, it is natural to compute curvature in this space as well. We set the ambient dimension, *n*, to be the number of PCs needed to explain 80% of the variance. The manifold dimension, *d*, for scRNAseq datasets is not well-defined and needs to be chosen heuristically. As a simple heuristic, we specified *d* as the number of PCs needed to explain 80% of the variance in the ambient space i.e. 64% of the original variance (we show later that our computations are relatively insensitive to the choice of *d*).

We considered three datasets. The first consists of *N* ≈ 10^4^ peripheral blood mononuclear cells (PBMCs) collected from a healthy human donor [42]. The second is a gastrulation dataset comprised of *N* ≈ 1.2 × 10^5^ cells pooled from 9 embryonic mice sacked at 6-hour intervals from embryonic day 6.5 to 8.5 [43]. The final sdataset is a benchmark in the field consisting of *N* ≈ 1.3 × 10^6^ brain cells pooled from 2 embryonic mice sacked at embryonic day 18 [44]. Refer to Figures S4A, S5A and S6A for cell type annotations for the three datasets.

The PBMC dataset is characteristic of the sample size of current scRNAseq data. The other two are larger than most scRNAseq datasets, and we included these to verify if geometric features seen in the first dataset can be reproduced for more densely sampled manifolds. For the PBMC, gastrulation and brain datasets, the ambient (manifold) dimensions were determined to be 8, 11 and 9 (3, 3 and 5) respectively, according to the aforementioned heuristic (see Methods Section 4.6.4). For all three datasets, the global length scale happened to be *L* ≈ 20 (see Methods Sections 4.3.5). As before, we picked *σ*_*h*_ for each dataset according to the distribution of GOF p-values (see Figures S4B, S5B and S6B; Methods Section 4.6.4).

We visualized the computed scalar curvatures on standard plots employed in the field (UMAP and t-SNE; shown in Figure 4A,D,G) and observed non-trivial scalar curvature for all three datasets. We found statistically significant correlations between the scalar curvature reported by each point and its kNN for *k* ≤ 250 (*ρ_Pearson_* = 0.58, 0.18 and 0.38 for the PBMC, gastrulation and brain datasets respectively at *k* = 250, *p* < 10^*−*6^; see Figures S4C, S5C and S6C), indicating that our algorithm yields scalar curvatures that vary continuously over the data manifolds. By plotting scalar curvatures against their standard errors, *σ*_*S*_, we verified that regions with non-zero scalar curvature are statistically significant (see Figure 4B,E,H). As a consistency check, we confirmed that the percentage of points with 95% CIs containing the scalar curvatures reported by their respective kNNs (i) decayed with increasing *k* for *k* ≤ 250, and (ii) was significantly larger than expected by chance (67%, 72% and 61% for the PBMC, gastrulation and brain datasets respectively at *k* = 250, *p* < 0.001; see Figures S4D, S5D and S6D; Methods Section 4.6.3.1).

**Figure 4:**
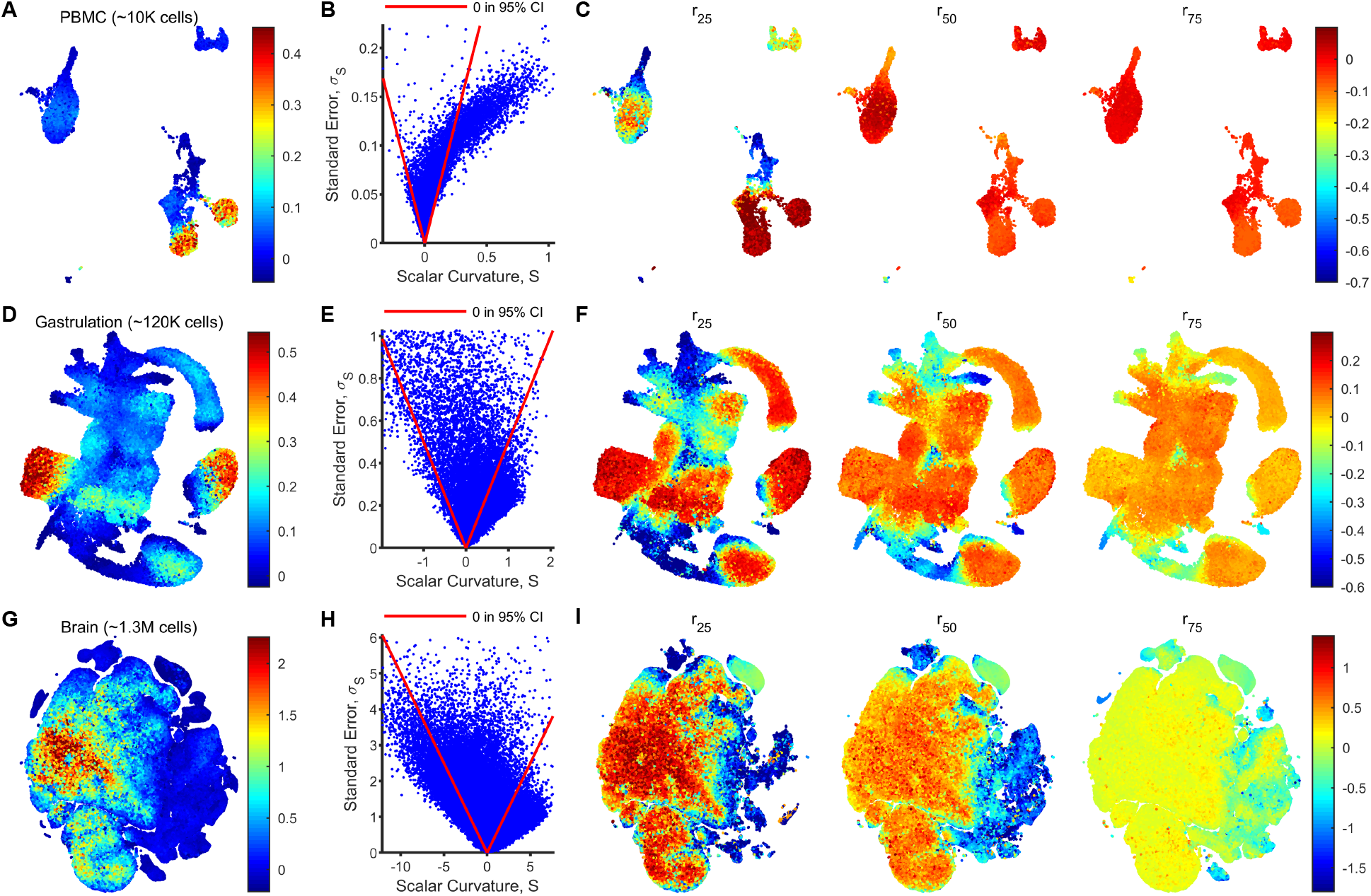
scRNAseq datasets have localized regions of non-zero scalar curvature. **(A)** Scalar curvatures were computed for a scRNAseq dataset with *N* ≈ 10^4^ peripheral blood mononuclear cells (PBMCs) collected from a healthy human donor. The ambient (*n*) and manifold (*d*) dimensions were specified to be 8 and 3 respectively and variable neighborhood sizes were chosen by setting *σ*_*h*_ (see Methods Section 4.6.4). The scalar curvatures are shown here overlaid onto UMAP coordinates, after smoothing the values over *k* = 250 nearest neighbors in the ambient space. **(B)** Scatter plot of (unsmoothed) scalar curvatures, *S*, and associated standard errors, *σ*_*S*_, for each datapoint in the PBMC dataset. Points enclosed by the red lines reported a 95% CI (*S* α 2*σ*_*S*_) including 0. **(C)** As in (A) but with scalar curvatures computed using a fixed neighborhood size, *r*, for all datapoints. The value of *r* was set to be the 25, 50, and 75-%ile values (left to right) of the neighborhood sizes used in (A) (see Figure S4E). Points for which a neighborhood of size *r* does not include enough neighbors for regression are not shown. **(D-F)** As in (A-C) for a mouse gastrulation dataset with *N* ≈ 1.2 × 10^5^, *d* = 3 and *n* = 11. **(G-I)** As in (A-C) for a mouse brain dataset with *N* ≈ 1.3 × 10^6^, *d* = 5 and *n* = 9, plotted on t-SNE coordinates.

To rule out the possibility that localization of non-zero scalar curvature in certain regions of the UMAP/t-SNE plots is an artifact caused by other features of the data that are also localized, we considered several factors. First, we plotted the GOF p-value at each point on UMAP/t-SNE coordinates and noted that poor GOFs were not localized on the data manifolds, let alone to regions of non-zero scalar curvature (see Figures S4B, S5B and S6B). Therefore, the computed scalar curvatures are not due to poor fits.

Next, we plotted the neighborhood size, *r*(*p*), used for fitting and observed that in some regions, non-zero scalar curvatures seemed to correspond to small *r* (see Figures S4E, S5E and S6E). Since *σ*_*h*_ is fixed, these regions necessarily have a larger number of neighbors *N*_*p*_(*r*) and are hence more dense (see Figures S4F, S5F and S6F). To rule out the possibility that the non-zero scalar curvatures were an artifact of smaller neighborhood size, we recomputed the scalar curvature at three fixed neighborhood sizes (see Figure 4C,F,I), corresponding to the 25, 50, and 75%-ile values of *r*(*p*) which arose from setting *σ*_*h*_ (see Figures S4E, S5E and S6E). In general, the scalar curvatures decreased in magnitude when neighborhood sizes increased. However, regions which had statistically significant non-zero scalar curvatures (zero falls outside of the 95% CI) using variable neighborhood sizes also had non-zero scalar curvatures for all three fixed neighborhood sizes. Additionally, statistically significant non-zero scalar curvature also emerged on other parts of the manifolds when using small fixed neighborhood sizes. These regions are therefore curved at small length scales but do not have a sufficient density of points to resolve curvature to the desired uncertainty *σ*_*h*_ (see Method Section 4.3.5). This is analogous to the image patch dataset for which we could resolve scalar curvatures of larger magnitude at a smaller length scale when the dataset was augmented with enough points to attain smaller neighborhood sizes for a fixed *σ*_*h*_.

We also checked how computed scalar curvatures changed with density in a toy model with zero scalar curvature. Importantly, we did not observe the artifactual appearance of statistically significant non-zero scalar curvature, for either variable neighborhood sizes chosen by the algorithm to achieve *σ*_*h*_, or for fixed neighborhood sizes (see Figure S2A; Methods Section 4.4.2.1). Taken together, although higher density allows us to resolve statistically significant non-zero scalar curvatures in scRNAseq data, these computed scalar curvatures are not an artifact of the smaller neighborhood sizes used in regions with higher density.

To ensure that the computed scalar curvatures were not sensitively dependent on the heuristically chosen manifold dimension, *d*, we also recomputed scalar curvatures for *d* − 1 and *d* + 1 and observed similar qualitative results (see Figures S4G, S5G and S6G). Lastly, we verified that the computed scalar curvatures were not correlated with the number of transcripts in each cell (see Figures S4H, S5H and S6H).

To confirm the robustness of our results to sampling, we randomly discarded *f* % of points in the ambient space determined for each dataset, and recomputed scalar curvatures using the same values of *n*, *d* and *r*(*p*) used for the original dataset. We found that a statistically significant percentage of downsampled points (82% for the PBMC dataset with *f* = 75, 78% for the gastrulation dataset with *f* = 75, and 76% for the brain dataset with *f* = 50; *p* < 0.001) had a 95% CI containing the scalar curvature reported by the same point for the original dataset (see Figures S4I, S5I and S6I; Methods Section 4.6.3.2). This suggests that if the datasets were more highly sampled, and scalar curvatures were recomputed using the same neighborhood sizes, they would be reliably contained within the currently reported 95% CIs. Unlike the two other datasets, the brain dataset could not be downsampled to *f* = 75 while still having at least 75% of points report 95% CIs containing the originally reported scalar curvatures, despite having the most points. This might be because the brain dataset has a larger manifold dimension according to our heuristic and therefore requires a greater number of terms, 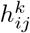, to be estimated in the Second Fundamental Form.

For the PBMC dataset, we additionally downsampled the single-cell count matrix by discarding *f* % of transcripts at random and preprocessing the same way. We recomputed scalar curvatures for this downsampled dataset with the same *n*, *d* and *r*(*p*) values used for the original dataset. Here too, we found that when *f* = 50 (*f* = 75), 70% (65%) of the downsampled points had a 95% CI containing the originally reported scalar curvature (*p* < 0.001, see Figure S4J; Methods Section 4.6.3.3). Therefore, the computed scalar curvature is robust to changes in capture efficiency and sequencing depth. Taken together, our computational analysis reveals non-trivial intrinsic geometry in scRNAseq data.

## 3 Discussion

In this study, we explored two approaches to computing the curvature of data manifolds using tools from twin branches of differential geometry. Despite the prevalence of the Laplace-Beltrami operator in geometric data analysis [14, 26, 27, 28, 29], an intrinsic approach to computing scalar curvature relying on this operator’s eigenvalues was determined to be infeasible for sample sizes of *N* ≈ 10^4^ typical of current scRNAseq datasets. Although methods such as MAGIC [45] and diffusion pseudotime [46] apply the Laplace-Beltrami operator to smooth scRNAseq data and infer cell differentiation trajectories respectively, using information intrinsic to the manifold, our results suggest that the embedding of the manifold in the ambient space provides valuable information necessary for estimating the intrinsic curvature. This observation is perhaps implicit in recent tools for estimating the Laplace-Beltrami operator, which first use moving local least-squares to approximate a surface, thereby incorporating information from the ambient space [29].

Certainly, we found that an extrinsic approach in which the embedding is retained, and curvature is determined by local quadratic fitting of datapoints in ambient coordinates, is feasible given the sample size and degree of noise in real-world datasets. To obtain the scalar curvature of data manifolds, our algorithm first computes the full Riemannian curvature tensor. For other applications, this tensor can be used to compute other geometric quantities, such as Ricci curvature, or may itself be of interest. More generally, we focused on intrinsic curvature because we were interested in geometric properties of the manifolds independent of their embeddings. However, the Second Fundamental Form used in our approach to compute the intrinsic curvature can be used to obtain all the information about the extrinsic curvature as well. Indeed, 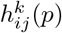 exactly quantifies the extent to which the manifold deviates in the *k*^th^ normal direction from the *ij*-tangent plane at point *p*.

A key limitation of our algorithm is that the manifold dimension must be specified by the user. We also assumed that the manifold dimension is the same at every point in a dataset. Extending the algorithm to determine the manifold dimension from the data itself, potentially in a position-dependent manner, may prove useful. In addition, there is no inherently correct length scale over which curvature should be computed for a data manifold. Our algorithm chooses a length scale that varies from one part of the data manifold to another according to the density of points, and is tuned to achieve a user-specified level of uncertainty in the computed curvature. For some applications, it might be more sensible to fix a desired length scale for computing the curvature.

As a demonstration of our algorithm, we computed the scalar curvature of image patches, and found that it was consistent with that of a Klein bottle. This observation further validates the claim by Carlsson et al. who showed that image patches have the topology of a Klein bottle [21]. Unlike the Klein bottle parameterization of image patches however, no definitive analytical form has been established for scRNAseq datasets. Recent work has suggested the use of hyperbolic geometry to model branching cell differentiation trajectories [47] and specific manifolds have been proposed to model reaction networks [48], which may be applicable to scRNAseq data. These proposed manifolds can be validated or improved using knowledge of the intrinsic geometry of scRNAseq datasets. Finally, incorporating information about curvature may provide a more principled approach for developing dimensionality reduction and visualization tools.

## 4 Methods

### 4.1 Differential Geometry of Theoretical Manifolds

Here we briefly discuss how to compute the scalar curvature of, and sample from, theoretical manifolds given a parameterization.

For a *d*-dimensional manifold, *M*, with intrinsic coordinates {*x*_1_,…, *x*_*d*_} and embedding in ℝ^*n*^ given by *f*(*x*_1_,…, *x*_*d*_), the metric is:

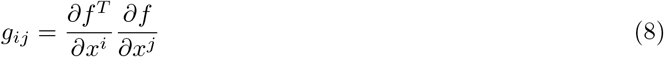

The scalar curvature of *M* can then be derived analytically in intrinsic coordinates in terms of the metric as

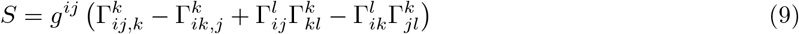

where the 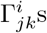 are Christoffel symbols given by

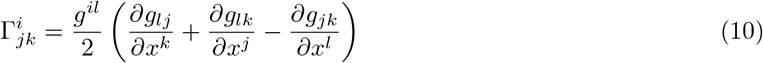

and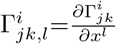

To draw points from *M* with *a*_*i*_ ≤ *x*_*i*_ ≤ *b*_*i*_ so that the embedded manifold is uniformly sampled in ℝ^*n*^, we use rejection sampling. For paired random variables *x* ∼ Uniform(*a, b*) and 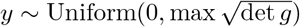,we retain *x* as a sample point if 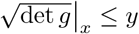.

### 4.2 Details of Intrinsic Approach to Curvature Estimation

Here we explain how we used Equations 2–4 on the simplest of toy manifolds, the noise-free 2-dimensional hollow unit sphere, 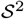, to obtain an estimate of the average scalar curvature. The true scalar curvature is *S*(*p*) = 2 ∀*p* ∈ *M*. For the remainder of this section, we adopt the convention that symbols with overbars are estimates of the corresponding unaccented quantities.

#### 4.2.1 Approach for 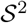

Our approach mirrors the treatment in [27], in which heat-traces are fit over various intervals [*x*_1_*, x*_2_] with *x*_1_ ≥ 0, to quadratic polynomials 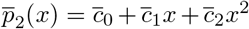 to estimate the geometric quantities in Equation 2.

Here, we constrained the form of 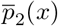 for fitting by assuming that (i) the manifold is boundary-less (so that 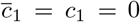 and the second boundary term for *c*_2_ vanishes), (ii) the volume is known (so that 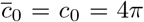), and (iii) the scalar curvature is constant (so that 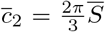), yielding 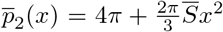. These are strong assumptions that will not hold for an arbitrary manifold, which already precludes this as a generic procedure. Nonetheless, we proceeded for 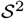 to see if even with this privileged information, the scalar curvature could be estimated accurately. We declared an estimate to be accurate on the interval [*x*_1_*, x*_2_] if 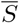 has error within ±0.5 i.e. 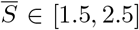. All quadratic fits were performed in MATLAB using the *lsqnonlin* function (‘StepTolerance’=1e-3, ‘FunctionTolerance’=1e-6).

First, we evaluated *z*_*m*_(*x*) using analytical eigenvalues for 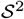 given by 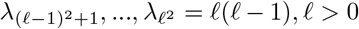, and let *D*_*m*_ be the collection of all intervals for which fits to 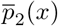 yielded accurate 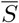. *D*_*m*_ corresponds to intervals where Equation 4 is accurate to our desired tolerance when the eigenvalues are known exactly.

Next, we uniformly sampled *N* = 10^4^ points from 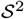 (see Figure 1A; Methods Section 4.4.1.1), estimated 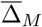 using the *random walk Graph Laplacian* with Gaussian kernel (see Equation 15 in Methods Section 4.2.5), and computed empirical eigenvalues, 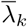, from 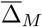. We selected *N* = 10^4^ as it is the same order of magnitude as the sample size of current scRNAseq experiments, and is sufficient to identify *M* as 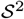 by eye (see Figure 1A). We verified if estimates 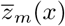, obtained by evaluating Equation 3 using 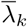, when fit as described above to 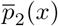 over intervals in *D*_*m*_, recapitulated the accurate 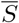 obtained using *z*_*m*_(*x*). We restricted our attention to *D*_*m*_ for calculations using empirical eigenvalues, since it is only over intervals in *D*_*m*_ that it is even theoretically possible to compute scalar curvature to the desired accuracy.

Below, we report our findings for different *m*.

#### 4.2.2 Infinite Series

We first applied this approach to the ideal case in Equation 3, where infinite analytical eigenvalues are available. We computed *z*_*∞*_(*x*) (shown as a black line in Figure S1A) and obtained 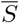 by fitting 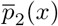 over various intervals as described above. Figure S1B shows that *D*_*∞*_ is comprised of intervals with 0 ≤ *x*_1_ < *x*_2_ ≲ 1.15. For *x*_2_ ≳ 1.15, errors from neglecting higher-order terms *o*(*x*^3^) in Equation 4 dominate. Since *z*_*m*_(*x*) converges from ∞, *x*_2_ ≲ 1.15 necessarily holds for any interval in *D*_*m*_ ∀*m*.

#### 4.2.3 Truncated Series

We next considered *z*_*m*_(*x*) for *m* < *N*, since in practice, we will only have access to as many eigenvalues as datapoints (*N*). We computed *z*_1000_(*x*) using Equation 3 (shown as a solid blue line in Figure S1A), and obtained 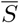 by fitting 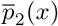 (see Figure S1C). Intervals in *D*_1000_ roughly satisfy 0.25 ≲ *x*_1_ < *x*_2_ ≲ 1.15. However, we found that 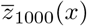 (shown as a dashed blue line in Figure S1A) deviated markedly from *z*_1000_(*x*) in the rough interval [0.1, 0.75], which has significant overlap with *D*_1000_. Consequently, when we fit 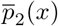 to 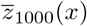 on *D*_1000_, the resulting 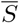 was not accurate for any interval in *D*_1000_ (see Figure S1D).

Note that this inaccuracy was not a consequence of not using all *N* available eigenvalues. While picking *m* = *N* would reduce the lower bound on valid intervals in *D*_*m*_ (since *z*_*m*_(*x*) converges from ∞), it is exactly for small *x*_1_ that 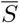 obtained from 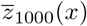 is already over-estimated as shown in Figure S1D. Since 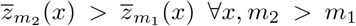, using a truncated series with a larger *m* would simply exaggerate the difference between *z*_*m*_(*x*) and 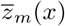 for small *x* and cause scalar curvatures estimated using the latter to be further over-estimated.

Following this line of thought, we reasoned that picking a fewer number of eigenvalues may ameliorate the issue. We selected *m* = 49 (instead of a round number like *m* = 50 so that all eigenvalues of a given multiplicity are included) and repeated this analysis for the same set of *N* = 10^4^ points. *z*_49_(*x*) is shown as a solid red line in Figure S1A and the intervals over which fits to 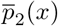 yield accurate 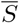, *D*_49_, are shown in Figure S1E. While 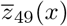 (shown as a dashed red line in Figure S1A) has a much smaller deviation from *z*_49_(*x*) than 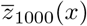 did from *z*_1000_(*x*), no estimate of 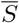 obtained from fits of 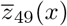 to 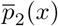 on *D*_49_ were sufficiently accurate once again (see Figure S1F).

#### 4.2.4 Eigenvalue Convergence

We refrained from reducing *m* further to improve agreement between 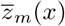 and *z*_*m*_(*x*) after noting that the size of the intervals in *D*_*m*_ shrink with *m*. Though we may have a better chance of computing accurate 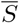 with 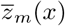 on *D*_*m*_ for smaller *m*, recall that in practice we will not have *D*_*m*_ available to us since the analytical eigenvalues will be unknown. Therefore, we simply shift the problem to one of choosing an interval that will yield an accurate 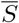, from a shrinking pool of intervals that could even theoretically yield an accurate estimate.

Instead, we compared the estimated 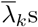 with their true values, *λ*_*k*_, and observed that the former consistently under-estimate the latter (see Figure S1H). Furthermore, we found that the fractional error grows with *k*, exceeding 60% for *k* = 37,…, 49. Therefore, 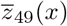 will only be accurate if *N* is large enough to limit the fractional error.

To determine the required tolerance on the fractional error, we constructed a truncated series analogous to Equation 3, but with eigenvalues interpolated between the analytical eigenvalues and the empirical eigenvalues determined for *N* = 10000, according to a parameter *f* :

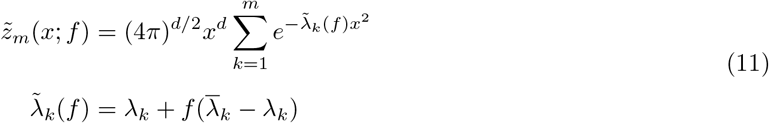

*f* signifies that the fractional error of the interpolated eigenvalues is reduced by 1− *f* relative to the empirical eigenvalues determined for *N* = 10000. We found that *f* ≤ 0.23 is needed so that 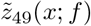 (shown as a green line in Figure S1A) fit to 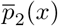 yields accurate 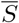 on half the intervals in *D*_49_ (see Figure S1G).

Given that the fractional error in estimating *λ*_37_*,…, λ*_49_ by 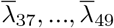 is 60% when *N* = 10000, how large does *N* have to be to reduce this fractional error to 60% × 0.23 ≈ 14%? A convergence rate for the fractional error is given in Theorem 1 of [30]. For 2-dimensional manifolds:

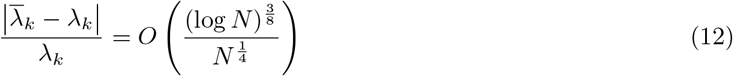

Assuming that the big-*O* bound is sharp at *N* = 10^4^ for *k* = 37*,…,* 49 (i.e. the prefactor is given by 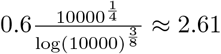 ≈ 2.61), we extrapolated that at least *N* = 10^7^ datapoints are needed to reduce the fractional error to 14% (see Figure S1H). Equation 12 also applies to empirical eigenvalues of 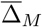 constructed from weighted kNN and *r*-neighborhood kernels instead of Gaussian kernels (see Methods Section 4.2.5). However, the prefactor in Equation 12 is actually worse for these estimators since their empirical eigenvalues have larger fractional errors at *N* = 10000 (see Figure S1H), so that even larger *N* would be required to attain the desired fractional error. Lastly, note that while we had analytical eigenvalues available with which to ascertain *m* = 49 as suitable, the naive approach of simply using all eigenvalues available (*m* = *N*), would require sample sizes that are even larger by several more orders of magnitude.

#### 4.2.5 Estimating the Laplace-Beltrami Operator from Data

For *N* points, {*X*_*i*_} ∈ ℝ^*n*^, sampled from *M*, we estimated Δ_*M*_ by normalizing the weight matrix *W* (see below) using the random walk normalization [30, 49]. 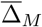 constructed using this normalization converges to Δ_*M*_ when samples are drawn uniformly from the embedding of *M* in ℝ^*n*^, as was done in our analysis.

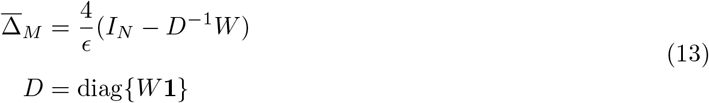

*I*_*N*_ is the *N* × *N* identity matrix, **1** ∈ ℝ^*N*^ is a vector of ones and the kernel width, *ϵ*, is set to match that used in Theorem 1 of [30]:

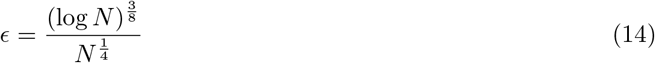

Throughout our analysis, we used *W* = *W*_*g*_, the weight matrix with entries given by a Gaussian kernel:

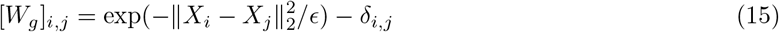

To check whether other estimators had more benign prefactors for eigenvalue convergence (see Figure S1H), we also considered the weighted kNN kernel, *W*_*kNN*_, and the *r*-neighborhood kernel, *W*_*r*_, with *r* = *ϵ* [50]:

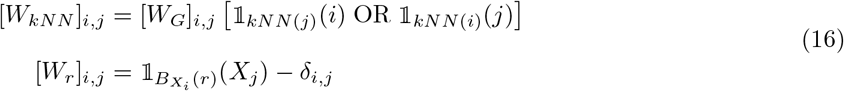

*kNN* (*i*) is the set of indices of the *k*-nearest neighbors of point *i* in ℝ^*n*^, 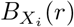 is the *n*-dimensional ball of radius *r* centred at *X*_*i*_, and 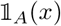 is the indicator function for *x* ∈ *A*.

### 4.3 Details of Extrinsic Approach to Curvature Estimation

#### 4.3.1 Quadratic Regression on Local Neighborhoods of Data

Here we describe the regression model for computing the coefficients of the Second Fundamental Form, 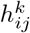, at a particular point *p*. As described in the main text, after performing PCA on a neighborhood of *N*_*p*_ points around *p* in ℝ^*n*^, each point in the neighborhood can be described in terms of *d* tangent coordinates, *t*_*i*_, and *n* − *d* normal coordinates, *n*_*k*_. We defer discussion of how the neighborhood is selected to Methods Section 4.3.2.

The *n*_*k*_s are treated as dependent variables that can be modelled as quadratic functions of the *t*_*i*_s, which are taken to be independent variables. See Equation 17 below. Linear terms are excluded since they ought to have zero coefficients in the tangent basis. Constant terms, *C*_*k*_, are included to account for affine shifts. Since 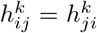 according to Equation 5, in practice we only consider *t*_*i*_*t*_*j*_ and 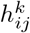 for *j* ≥ *i* so that **t** and **h** in Equation 17 have linearly independent columns, though we write the full form here for simplicity.

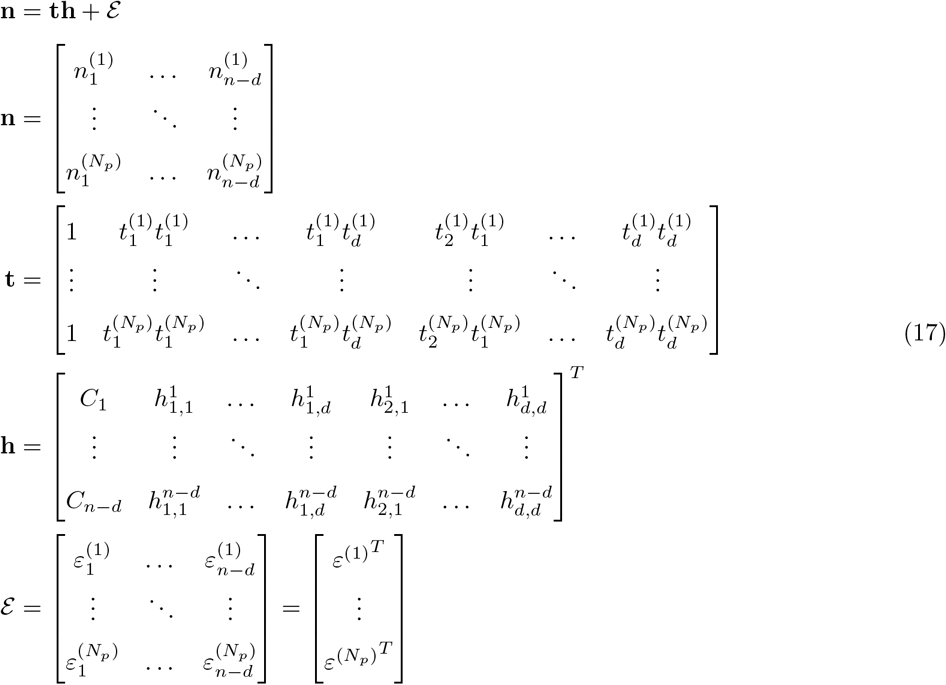

Regression yields the following least-squares solution:

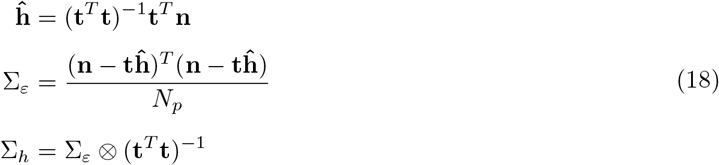

where **ĥ** is the matrix of estimates of the Second Fundamental Form, Σ_*∊*_ is the estimated covariance structure of the residuals so that 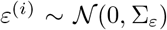, and Σ_*h*_ is the covariance matrix for **ĥ**. ⊗ denotes the Kronecker product. We used the *mvregress* function in MATLAB to perform this regression in our code.

When datapoints are sampled exactly from an analytical manifold, Σ_*∊*_ measures the contribution of higher-order terms. In the limit of infinite sampling and infinitesimally small neighborhoods, Σ_*∊*_ → 0. When observational noise is present (discussed in Methods Section 4.4.2.2), Σ_*∊*_ also depends on the magnitude of the noise (*σ* in Equation 28).

#### 4.3.2 Selecting Local Neighborhoods for Regression

Here we describe the procedure for selecting a neighborhood around each point *p* for computing the Second Fundamental Form. We adopt the simplest approach of selecting the neighborhood to be a ball of radius *r* centred at *p*, *B*_*p*_(*r*).

If *r*(*p*) is not specified, we set it according to statistical rather than geometric principles, since the geometry of the manifold may be non-trivial and unknown *a priori*. Specifically, we set *r*(*p*) so that the elements in the covariance matrix, Σ_*h*_, are upper-bounded by 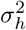, the square of the specified target uncertainty. The largest elements in Σ_*h*_ are the variance terms on the main diagonal, corresponding to the squares of the standard errors, 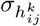, for the coefficients 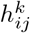. By inspection of Equation 18:

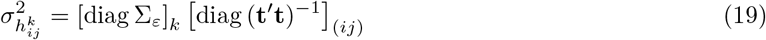

where [diag Σ_*∊*_]_*k*_ is the diagonal entry of Σ_*∊*_ corresponding to the *k*^th^ normal direction and [diag (**t**′**t**)^*−*1^]_(*ij*)_ is the diagonal entry in (**t**′**t**)^*−*1^ for which the corresponding entry in **t**′**t** is 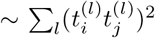. Increasing *r*(*p*) monotonically increases both *N*_*p*_(*r*), the number of points in *B*_*p*_(*r*), and the average magnitude of elements in **t**, both of which reduce 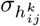.

To avoid sweeping *r*(*p*) to find the minimum value such that 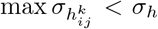, which is computationally expensive, for each point we instead model the dependence of *N*_*p*_(*r*) on *r* as

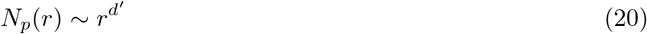

so that

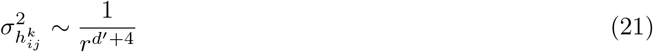

To determine *d*′, *N*_*p*_(*r*) is counted at 10 log-spaced distances, *r*_*i*_, and a line is fit to the (log *r*_*i*_, log *N*_*p*_(*r*_*i*_)) pairs for *i* ∈ {2, …, 8}. *r*_1_ is set to be the distance from *p* to the 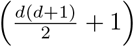-closest point to *p* (the minimum number of points needed for regression). *r*_10_ is set to be the distance from *p* to the furthest point from *p*. To solve for *r*, we first guess *r*_*g*_ = *r*_1_, perform regression on the set of points in *B*_*p*_(*r*_*g*_) and assign 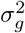 to be the largest diagonal entry in Σ_*h*_. If 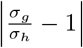 is within a desired tolerance, we set *r* = *r*_*g*_, or else we update *r*_*g*_ as shown and iterate to convergence.

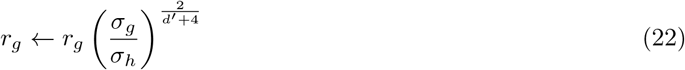

For large datasets, we speed up computation by only selecting *r* in this manner for a subset of *N*_*calib*_ ≤ *N* randomly selected calibration points. All datapoints in the Voronoi cell of each calibration point are then assigned the same *r* as the calibration point. Unless otherwise specified *N*_*calib*_ = *N*.

#### 4.3.3 Goodness-of-Fit Test for Quadratic Regression

For a fixed density of points, there is a fundamental trade-off between reducing uncertainty in the 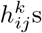 and the validity of approximating local neighborhoods with quadratic fits. To reduce *σ*_*h*_, more points must be included in the fit, but a larger neighborhood may not be well-modelled by only quadratic terms. Conversely, 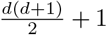 points are sufficient to perform the regression, but there is then large uncertainty in the estimate of 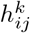. Since our approach is to choose a neighborhood size to achieve a target *σ*_*h*_, we include a companion goodness-of-fit (GOF) statistic measuring how well the neighborhood is fit by a quadratic. Namely, we use Mardia’s test on the residuals from regression (*∊*^(*i*)^ in Equation 17), which yields a p-value for the null hypothesis that the residuals are normally distributed [51].

When the p-values are small, the quadratic regression model is unlikely to be valid. In this case, curvatures computed using the resulting 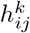 may be suspect regardless of the tightness of the errorbars, and the user may want to consider increasing *σ*_*h*_ to reduce the neighborhood size. However, the poor GOF may not be of concern if the length scale of interest is larger than the fluctuations in the manifold which give rise to the non-Gaussian residuals (see Methods Section 4.3.5). Note that Mardia’s test is relatively weak since it may yield false negatives for heteroskedastic residuals. This GOF measure is therefore only provided as a computationally cheap consistency check. Ideally, the density of sampled points is sufficiently high to (i) permit small *σ*_*h*_ and (ii) produce GOF p-values that are uniformly distributed (consistent with the null model) and spatially uncorrelated.

#### 4.3.4 Standard Error and Bias of Scalar Curvature Estimate

Here we discuss how we compute the standard error, *σ*_*S*_, of the estimate for *S* and note sources of estimator bias. Since the Riemannian curvature tensor in Equation 6 is a bilinear form and the tensor contraction in Equation 7 is a straightforward sum, *σ*_*S*_ can be computed using simple error propagation formulas in terms of the uncertainties from regression. Specifically, the standard error we report is the first-order approximation to the second moment of a function of random variables:

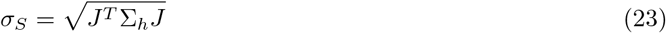

where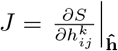.

It is important to note that our estimate for *S* is biased and not normally distributed. First, the 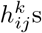 are only normally distributed when the residuals (*∊*^(*i*)^ in Equation 17) themselves are normally distributed. Second, even when the 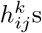 are normally distributed, our estimate of *S* will not be due to its bilinear dependence on 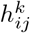. Lastly, estimates for *S* can be biased in a manifold-dependent and even position-dependent way. For instance, the analytical scalar curvature of 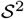 embedded in ℝ^3^ is given by 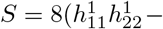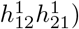, with 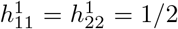 and 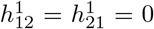. Numerically however, the symmetric off-diagonal terms will never be exactly 0 so *S* will be systematically under-estimated. This is apparent in the left tail of the blue histogram in Figure 2I. In our experience, adding isotropic noise of small magnitude tends to remove the skew, presumably because then the residuals more closely match the regression assumptions (see for example Figure S2B, where the left tail disappears for *σ* = 0.001). Furthermore, in our examples, we observed that computed scalar curvatures were less biased when the ambient and/or manifold dimensions were large. We speculate that this is because the increased number of terms (with alternating signs) in Equations 6 and 7 leads to cancellation of errors, which is likely why the accuracy of computed scalar curvatures was higher for 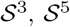 and 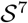 than 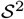, and the distribution of scalar curvatures less skewed (see Figure 2I).

#### 4.3.5 Note on Length Scales

Here we make three remarks regarding length scales relevant both for considering curvature theoretically and for applying our algorithm.

First, note that scalar curvature has units of inverse length squared. Therefore, scaling all the coordinates of the points on a manifold by a factor *L*, changes the scalar curvature at all points by *L*^−2^. Thus, it is always important to contextualize the scalar curvature in terms of the global length scale associated with the manifold. For example, the scalar curvature of 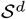 with radius *R* is 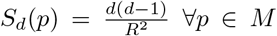 (here *L* = *R*). In the case of the toy models shown in Figure 2, the global length scale is *L* ≈ 1 (see Methods Section 4.4.1). For the image patch dataset, a normalization is applied which places all patches on 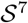 (see Methods Section 4.5.2), so that the global scale is again *L* = 1. For scRNAseq data, we computed scalar curvature on the datapoints after preprocessing (see Methods Section 4.6.1), without imposing any additional scaling correction to achieve a standardized global length scale. Since other custom analyses also use these same boilerplate preprocessing steps, computing scalar curvatures in the context of the global length scale of the preprocessed data is sensible. For all three scRNAseq datasets, the global length scale happened to be *L* ≈ 20 (see Methods Section 4.6.4).

Second, since 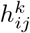 is a dimension-ful quantity (which scales as *L^−^*^1^), to keep the ratio of *σ*_*S*_ to *S* fixed when all coordinates are scaled by *L*, *σ*_*h*_ needs to be scaled by *L^−^*^1^.

Lastly, we note that our choice of *σ*_*h*_ sets local length scales that are statistically rather than geometrically informed: neighborhoods are chosen to upper bound the uncertainty in estimates obtained from regression. This length scale can also be understood in terms of a bias-variance trade-off. Large length scales reduce variance but may introduce a bias if the resulting neighborhoods are larger than features on the manifold. This manifests as poor GOFs and can be corrected by finer sampling. However, for manifolds with features at different length scales (such as a golf ball, which can be treated as dimples superimposed on 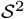), neigh-borhoods chosen by this heuristic can also be much smaller than the feature of interest, so that fine-scale curvature fluctuations are detected (dimples) while coarser features are neglected 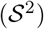. Regardless, we default to this statistical approach because in general, the length scale of relevant features on a data manifold will not be uniform across the manifold or known *a priori*. However, we also provide the ability to manually set position-dependent *r*(*p*) in the software to facilitate *ad hoc* computation of curvatures at any length scale of interest.

### 4.4 Details of Toy Manifold Curvature Computations

#### 4.4.1 Analytical Forms

Here we provide analytical forms for the toy manifolds shown in Figures 2 and S2.

##### 4.4.1.1 Hypersphere

The *d*-dimensional unit hypersphere, 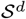, has intrinsic coordinates *θ*_1_ ∈ [0, 2*π*], 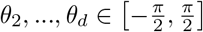 and ambient coordinates in ℝ^*d*+1^ given by:

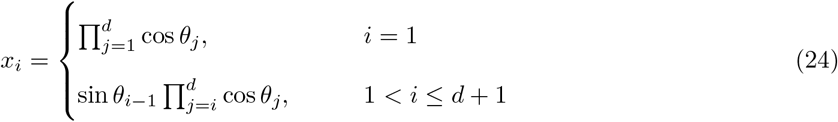

Using the relations in Methods Sections 4.1, the scalar curvature is given by *S*_*d*_(*p*) = *d*(*d* − 1) ∀*p* ∈ *M*. To draw uniform samples from 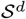, instead of applying rejection sampling on these intrinsic coordinates as described in Methods Section 4.1, it is more straightforward to let 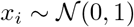 and scale the resulting vector (*x*_1_,…, *x*_*d*+1_) to have unit norm.

##### 4.4.1.2 One-Sheet Hyperboloid

The one-sheet hyperboloid, 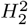, has intrinsic coordinates *θ* ∈ [0, 2*π*], *u* ∈ ℝ and ambient coordinates in ℝ^3^ given by:

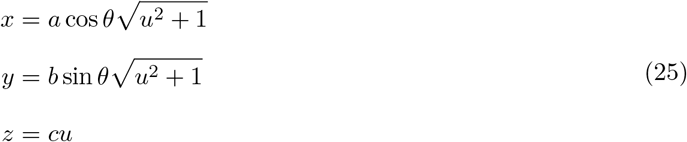

For Figure 2E,F, we used *a* = *b* = 2 and *c* = 1. Using the relations in Methods Sections 4.1, the scalar curvature is given by 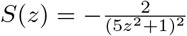. To avoid edge effects in the *z*-direction, we constrained *u* ∈ [−2, 2], and sampled points as described in Methods Section 4.1 until a subset of *N* = 10^4^ had *u* ∈ [−1, 1]. Scalar curvature was computed and visualized for these *N* = 10^4^ points.

##### 4.4.1.3 Ring Torus

The 2-dimensional ring torus, *T*^2^, has intrinsic coordinates *θ, ϕ* ∈ [0, 2*π*] and ambient coordinates in ℝ^3^ given by:

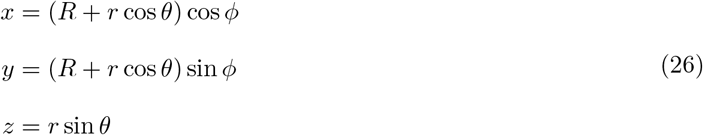

For Figure 2G,H, we used *R* = 2.5 and *r* = 0.5. Using the relations in Methods Sections 4.1, the scalar curvature is given by 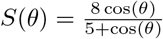

##### 4.4.1.4 Hypercube

The *m*-dimensional cube of side length *r*, 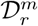, has intrinsic coordinates *z*_1_,…, *z*_*m*_ ∈ [−*r/*2*, r/*2], and ambient coordinates in ℝ^*n*^ for *n* ≥ *m* given by:

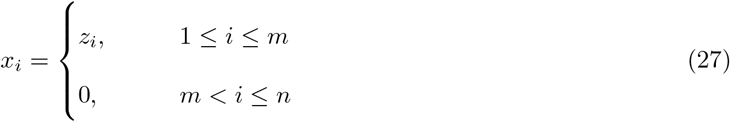

Using the relations in Methods Sections 4.1, the scalar curvature is given by *S*(*p*) = 0 ∀*p* ∈ *M*.

#### 4.4.2 Practical Issues for Curvature Estimation on Real-World Datasets

For real-world data, small sample size is only one of the potential confounders for accurately estimating curvature. Here, we report how our algorithm fares when four other real-world confounders are applied to toy manifolds: non-uniform sampling, observational noise, large ambient dimension *n*, and uncertainty in the manifold dimension *d*.

##### 4.4.2.1 Non-Uniform Sampling

We expect our approach to handle non-uniform sampling of the manifold gracefully: smaller (larger) neighborhoods will be used on densely (sparsely) sampled portions of the manifold to encapsulate the number of points needed to achieve *σ*_*h*_. To computationally verify the robustness of our tool to non-uniform sampling, we constructed a toy model to roughly match the (*n*, *d*, *L*) parameters for the scRNAseq datasets explored in the paper, for which non-zero scalar curvatures seemed to appear at smaller length scales/higher densities. Specifically, we wanted to verify that non-zero scalar curvatures do not appear artifactually at specific length scales due to sharp changes in the local density of points sampled from a flat manifold. To this end, we formed a dataset with a sparse periphery and dense core by uniformly sampling *N*_1_ = 10^4^ points from 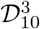 to establish a background density equal to 10 points per unit volume, and *N*_2_ = 10^3^ points from 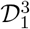 to create a core density roughly equal to 10^3^ points per unit volume (see Methods Section 4.4.1.4). We embedded these points in ℝ^11^ by adding isotropic Gaussian noise with *σ* = 0.01 to the eight normal directions, for all datapoints.

We computed scalar curvature on this dataset for a fixed *σ*_*h*_ (see Methods Section 4.4.3) and found no significant deviation from the true value of zero in either the sparse or dense regions (see Figure S2A). We next computed scalar curvatures at three fixed length scales corresponding to the 5, 50, and 95%-ile *r* values obtained using the specified *σ*_*h*_ (*r* = 0.54, 0.90 and 1.22 respectively) and again saw no deviation from zero scalar curvature for points in either the sparse or dense region (see Figure S2A). We repeated this analysis for *N*_2_ = 10^4^ and again saw no deviation from zero scalar curvature, regardless of whether variable neighborhood sizes or fixed length scales (*r* = 0.37, 0.63 and 1.42 corresponding to the same percentiles) were used (see Figure S2A).

##### 4.4.2.2 Observational Noise

Every ambient coordinate can be considered a measured observable with its own observational noise. Assuming each observable is distorted by independent, isotropic Gaussian noise with variance *σ*^2^ (sometimes referred to as *convolutional* noise [37]), datapoints *X* ∈ ℝ^*n*^ sampled from an embedded manifold *M* are modelled by:

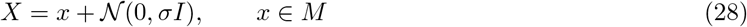

To study the sensitivity of our algorithm to noise, we uniformly sampled *N* = 10^4^ datapoints from 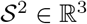, added convolutional noise with *σ* ranging over several orders of magnitude, and estimated scalar curvatures using a fixed *σ*_*h*_ (see Methods Section 4.4.3). For small *σ*, the distribution of scalar curvatures was centred on the true value of 2, but once *σ* became large (≈ 10% of 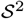’s radius), the estimated scalar curvatures approached 0 (see Figure S2B). Noise in the regression context does not change the expectation value of any estimated parameter. The apparent flattening that is observed therefore indicates that *X* (obtained from convoluting *M*), has a geometry that is not trivially related to *M*. Certainly for *σ* ≈ 1, *X* does not even preserve the topology of *M* as 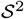. From a practical perspective, it suffices to say that small convolutional noise can be handled by simple quadratic regression, while large convolutional noise obfuscates the original manifold.

These observations are consistent with literature defining a manifold’s *reach* [34, 35], a noise scale beyond which noisy samples cannot be uniquely associated to a point on the noise-free manifold. When *σ* exceeds the manifold’s reach, the relationship between the empirical density of sampled points and the original manifold is non-trivial even for a relatively forgiving model of manifold-orthogonal noise. The *ridge* manifold [36, 37] of an empirical density has also been defined as an alternative to the unwieldy task of deconvoluting noisy samples to recover a noise-free manifold. This definition avoids the notion of a noise-free manifold altogether and instead defines manifolds as ridges, contours along which the empirical density of points is maximized.

##### 4.4.2.3 Large Ambient Dimension

A high-dimensional dataset may have an ambient space comprised of tens of thousands of observables, i.e. *n* is very large. Meanwhile, the underlying manifold dimension, *d*, may be small. Since convolutional noise occurs in *n* dimensions, will a low-dimensional manifold still be discernable?

To explore this, we uniformly sampled *N* = 10^4^ datapoints from 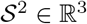, embedded these points in ℝ^*n*^ for a range of *n* up to 100, and added convolutional noise of magnitude *σ* = 0.01, 0.03, and 0.05 in the *n*-dimensional ambient space. We computed curvatures for all combinations of *n* and *σ* using a fixed *σ*_*h*_ (see Methods Section 4.4.3). As *n* or *σ* increased, the algorithmically chosen neighborhood sizes, *r*(*p*), expanded to include enough datapoints to maintain the desired *σ*_*h*_. The distribution of estimated scalar curvatures (shown in Figure S2C) is centred on the true value of 2 for *n* < 80 and *σ* ≤ 0.05.

However, we observed that *r* was far less sensitive to changes in *n* than changes in *σ*. For example, exploding *n* from 3 to 100 at *σ* = 0.01 and tripling *σ* from 0.01 to 0.03 at *n* = 3 required a comparable increase in *r* (see Figure S2C). Therefore, consistent with the results of Methods Section 4.4.2.2, as long as the noise scale *σ* is small, a large ambient dimension *n* is not a confounder. Practically however, to shorten computational overhead and avoid the large-*n*-and-*σ* case, it is still helpful to reduce the ambient dimension by projecting datapoints to an affine subspace containing the manifold (e.g. by PCA). Such a transformation does not change the intrinsic curvature.

##### 4.4.2.4 Choice of Manifold Dimension

The last practical consideration is accurate selection of the manifold dimension, *d*, which we have so far assumed to be known. There is no consensus on the definition of *d* for a dataset, so various disciplines have devised different heuristics to determine *d* in a data-driven fashion [52]. From the regression perspective, any *d >* 0 corresponds to a well-defined regression problem. The choice of *d* merely determines how local coordinates are partitioned into independent (tangent) and dependent (normal) variables. However, in our algorithm we noticed that some choices of *d* result in excessively large *r*(*p*) for a fixed *σ*_*h*_. We explored this further using two toy manifolds and discovered a signature indicating that the specified manifold dimension may be incorrect.

The manifolds considered were 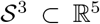 convoluted by isotropic Gaussian noise with *σ* = 0.01 and 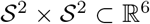, for which *d*\*, the true manifold dimension, is *d*\* = 3 and *d*\* = 4 respectively. We uniformly sampled *N* = 10^4^ points from each manifold and estimated scalar curvatures by holding *σ*_*h*_ fixed for different *d* (see Methods Section 4.4.3). For both manifolds, the average neighborhood size, *r*, was much larger for *d* > *d*\* and *d* < *d*\*, than for *d* = *d*\* (see Figure S2D). In the case of 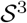, for *d* < *d*\*, the average neighborhood size was even larger than the global length scale, *L*, of the manifold. Since neighborhood sizes are chosen to achieve a target *σ*_*h*_, manually decreasing *r*(*p*) is counter-productive and simply increases the uncertainty from regression above *σ*_*h*_.

The large neighborhood sizes that emerged for both *d* > *d*\* and *d* < *d*\* can be understood in terms of the mis-assignment of normal vectors to the tangent space, or vice versa. According to Equation 19, 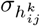 increases with large variation in the normal direction ([diag Σ_*∊*_]_*k*_), or with small variation in the tangent direction ([diag (**t′t**)^−1^]_(*ij*)_). When we choose *d* > *d*\*, we mis-attribute a normal direction with small variation [diag Σ_*∊*_]_*k*_ as an independent variable, whereas variation along the true tangent space is ≫ [diag Σ_*∊*_]_*k*_. *r* must therefore be increased to compensate for the lack of variation along this direction mis-classified as tangent. When *d* < *d*\*, we have spuriously assigned a tangent direction with large variation to be a normal direction. Since this spurious normal coordinate cannot be well-approximated as a function of tangent coordinates from which it is linearly independent, the perceived noise scale ([diag Σ_*∊*_]_*k*_) is exaggerated so that a larger neighborhood is needed to attain *σ*_*h*_.

This suggests a crude, operational definition of what constitutes an incorrect choice of *d*. When 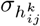 is large relative to the uncertainty in other coefficients, there is either too little variation along the *i*^th^ and *j*^th^ tangent directions, or too much variation along the *k*^th^ normal direction. In the former case, the *i*^th^ or *j*^th^ tangent direction might be more appropriately classified as a normal direction (*d* is too large and should be decreased), while in the latter case, the *k*^th^ normal direction might be more appropriately classified as a tangent direction (*d* is too small and should be increased). When this criterion is applied point-wise, there may be a different acceptable choice of *d* for different parts of the manifold. When this criterion is generalized over the entire manifold, a *σ*_*h*_ yielding a flat distribution of GOF p-values when the manifold dimension is specified to be *d* will also yield a flat distribution for *d* + 1 but not necessarily for *d* − 1: if residuals in *n* − *d* dimensions are well-modelled by a multivariate Gaussian, so too will residuals in *n* − *d* − 1 dimensions, but not necessarily residuals in *n* − *d* + 1 dimensions (see Figure S2D). Our observations are consistent with manifolds in literature with multiple possible manifold dimensions (like the helix manifold in [36]), and which could generally arise from non-isotropic noise or non-uniform sampling.

#### 4.4.3 Parameters for Curvature Estimation

For each manifold in Figure 2, we chose *σ*_*h*_ so that the fraction of points with GOF p-value ≤ *α* = 0.05 most closely matched the null model of normally distributed residuals consistent with neighborhood sizes well-approximated by quadratic regression (see Section 4.3.3). *σ*_*h*_ = (0.017, 0.020, 0.028, 0.055, 0.022, 0.030) for 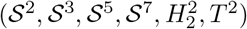 resulted in (7.4, 3.5, 1.4, 2.8, 7.4, 4.0)% of points having GOF p-values ≤ *α* = 0.05. Theoretically, 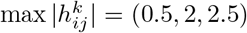 for 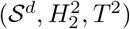 so our choices for *σ*_*h*_ result in small fractional errors in all cases.

For Figure S2A, we set *σ*_*h*_ = (0.02, 0.01) for *N*_2_ = (10^3^, 10^4^) respectively which resulted in (1.6, 2.5)% of points having GOF p-values ≤ *α* = 0.05. For all other panels in Figure S2, where we were interested in ascertaining the sensitivity to different confounders, instead of minimizing uncertainty per se, we used a fixed value of *σ*_*h*_ = 0.05. This choice resulted in neighborhoods small enough to be well-approximated by quadratic regression, manifesting as a roughly uniform distribution of GOF p-values in all cases.

### 4.5 Details of Image Patch Dataset and Klein Bottle Manifolds

#### 4.5.1 Notation and Preliminaries

First we introduce some notation needed to describe the image patch dataset. We refer readers to [21, 40] for a more detailed exposition. Let *P* be the space of all bivariate polynomials *p* : ℝ × ℝ → ℝ with *p* ∈ *P*, *h* : *P* → ℝ^9^ the vectorization operator given by *h*(*p*) = [*p*(−1, 1)*, p*(−1, 0)*, p*(−1, −1)*, p*(0, 1)*, p*(0, 0), *p*(0, −1), *p*(1, 1), *p*(1, 0), *p*(1, −1)]^*T*^, 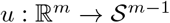 the normalization operator given by 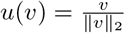, and *c* : ℝ^9^ → ℝ^8^ the projection operator given by *c*(*y*) = Λ*A^T^ y*, where *A* = [**e**_**1**_ … **e**_**8**_], 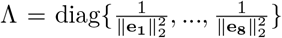, and {**e**_**1**_} are vectorized basis vectors for the 2-dimensional discrete cosine transform (DCT) applied to 3×3 patches:

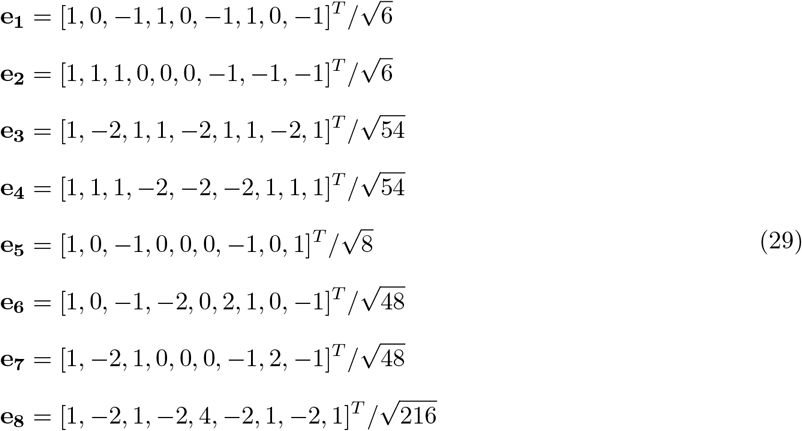

By inspection, **e**_**1**_ is the basis vector for patches with horizontal stripes and linear gradients, **e**_**2**_ for patches with vertical stripes and linear gradients, **e**_**3**_ for patches with horizontal stripes and quadratic gradients, **e**_**4**_ for patches with vertical stripes and quadratic gradients, and **e**_**5**_ for diagonally-oriented patches with quadratic gradients. All the patches produced by the embedding *k*^0^ in Equation 31 below and visualized in Figure 3B can be written as a linear combination of these 5 basis vectors. Next, note that the components in each **e**_**i**_ sum to 0, so that the projection operator, *c*, additionally serves to remove the mean. Finally, observe that the vector norm formed under *D* = *A*Λ^2^*A^T^* (referred to hereafter as the D-norm following [40]) measures the contrast in a 3×3 patch since

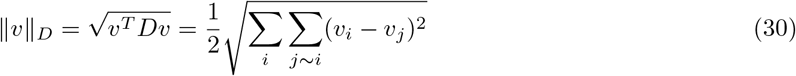

where *j* ∼ *i* refers to all vertical and horizontal neighbors, *j*, of a pixel *i* in the preimage of *v* under *h*. The **e**_**i**_ are normalized so that ||**e**_**i**_||_*D*_ = 1.

#### 4.5.2 Image Dataset

We used the same *van Hateren* IML dataset [41] consisting of 4167 greyscale images of size 1532×1020 pixels studied by Carlsson et al. in [21] and followed the same preprocessing steps used there. In short, we applied a *log1p* transformation to all pixel values and randomly sampled 5 × 10^3^ (possibly overlapping) 3×3 patches from each image. We indexed the pixels in each patch using standard Cartesian coordinates with the middle pixel as the origin, so that log-transformed pixel values are given by *p*(*x, y*), *x* ∈ {−1, 0, 1}, *y* ∈ {−1, 0, 1}. We then applied *h* to vectorize each patch *p*, and retained the high-contrast patches comprising the top quintile of D-norms for each image, resulting in *N* ≈ 4.2 × 10^6^ datapoints. Next, we normalized these high-contrast vectorized patches using the composition *u* ◦ *c*, resulting in a set of datapoints on 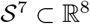. We determined the density of these datapoints in ℝ^8^ using the kNN density estimator with *k* = 100, and retained the densest decile, which yielded *N* ≈ 4.2 × 10^5^ datapoints. This dense subset of high-contrast normalized patches was found using topological data analysis in [21] to be a Klein bottle, 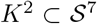, and is studied in Figures 3D,I and S3B.

To generate the augmented image patch dataset used in Figures 3J and S3E,F, we first considered all *N* ≈ 1.3 × 10^9^ vectorized high-contrast patches in the *van Hateren* IML dataset using the same procedure described above (each of the 4167 images yields 1530 × 1018 patches, of which the top 20% by D-norm are retained per image). These were normalized by *u* ◦ *c* as before to place them on 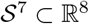. We again wanted to retain the densest decile of points, since only these have the topology of a Klein bottle. Mirroring the approach in [21] where the *k* used in the kNN estimator was scaled with sample size, *k* = 10^2^ used for *N* ≈ 4.2 × 10^6^ corresponds to 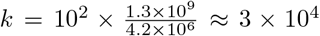 for *N* ≈ 1.3 × 10^9^. Computing *k* ≈ 3 × 10^4^ neighbors for all *N* ≈ 1.3 × 10^9^ points is prohibitive however. To determine a reasonable smaller value of *k*, we randomly selected 2 × 10^4^ points from the set of *N* ≈ 1.3 × 10^9^ on which to compare estimators and found that 90% of points in the densest decile as computed with *k* = 3 × 10^4^ also appeared in the densest decile computed using *k* = 6 × 10^2^. We therefore used the latter value for density estimation and retained the *N* ≈ 1.3 × 10^8^ datapoints comprising the densest decile.

#### 4.5.3 Parametric Family of Klein Bottle Embeddings

Let *θ, ϕ* ∈ [0, 2*π*]. Bivariate polynomials parameterized by (*θ, ϕ*), *k*_*θ,ϕ*_ ∈ *K*_*θ,ϕ*_ ⊂ *P*, that satisfy *k*_*θ,ϕ*_ = *k*_*θ*+*π*,2*π*−*ϕ*_ form a Klein bottle, *K*^2^: the (*θ, ϕ*) ∼ (*θ* + *π,* 2*π* − *ϕ*) similarity relation results in edges being glued together in the manner definitional of a Klein bottle’s topology (shown in Figure 3B). The candidate Klein bottle embedding supplied in [21] to model image patch data satisfies the similarity relation ∀*x, y*:

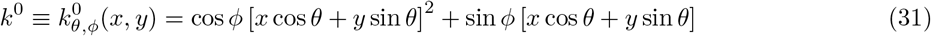

Note that any *k*_*θ,ϕ*_ ∈ *K*_*θ,ϕ*_ can be decomposed as:

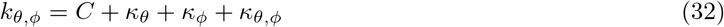

where *κ*_*θ*_ = *κ*_*θ*+*π*_, *κ*_*ϕ*_ = *κ*_2*π*−*ϕ*_ and *κ*_*θ,ϕ*_ = *κ*_*θ*+*π,*2*π*−*ϕ*_. The first three terms can be understood as constant, *θ*-dependent and *ϕ*-dependent phases respectively.

We sought an embedding of the Klein bottle for which the sum of Euclidean distances from each image patch to its closest point on the embedding is minimized. To accomplish this, we constructed a parametric family of models for each of the four terms in Equation 32. The first three of these are most conveniently expressed directly in the DCT basis.

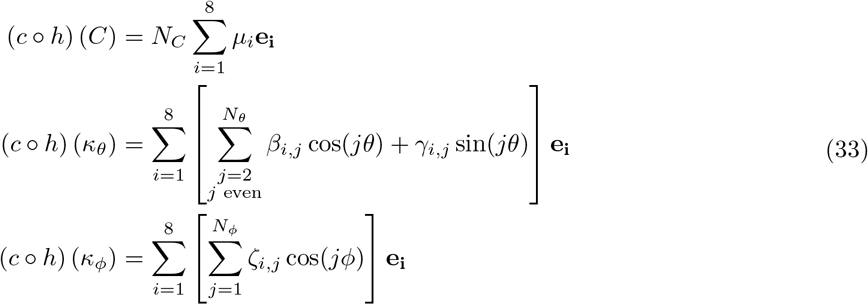

*N*_*C*_ is a Boolean variable, and *N*_*θ*_ and *N*_*ϕ*_ control the number of terms in the inner sum for (*c* ◦ *h*) (*κ*_*θ*_) and (*c* ◦ *h*) (*κ*_*ϕ*_) respectively. The expression for (*c* ◦ *h*) (*κ*_*θ*_) only includes even coefficients for *θ* so that the similarity relation (*θ*) ∼ (*θ* + *π*) is satisfied. The expression for (*c* ◦ *h*) (*κ*_*ϕ*_) only includes cosine terms so that the similarity relation (*ϕ*) ∼ (2*π* − *ϕ*) is satisfied.

For *κ*_*θ,ϕ*_, we refrained from writing a Fourier series-like expansion because we wanted to preserve the interpretation of *θ* and *ϕ* as parameters controlling the orientation and gradient respectively [21]. Instead, we devised the following form, which we motivate further below:

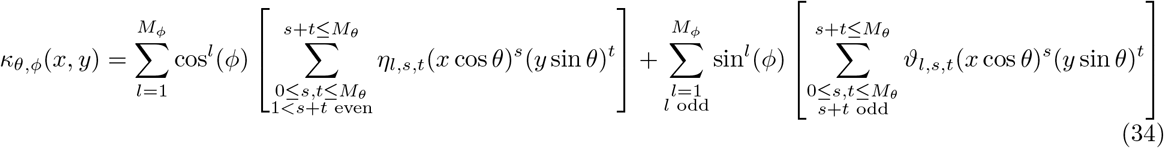

The expression for (*c* ◦ *h*) (*κ*_*θ,ϕ*_) is unwieldy but we note the following identities for monomials of the form *q*(*x, y*) = *x*^*s*^*y*^*t*^, which can then be applied term-wise to *κ*_*θ,ϕ*_.

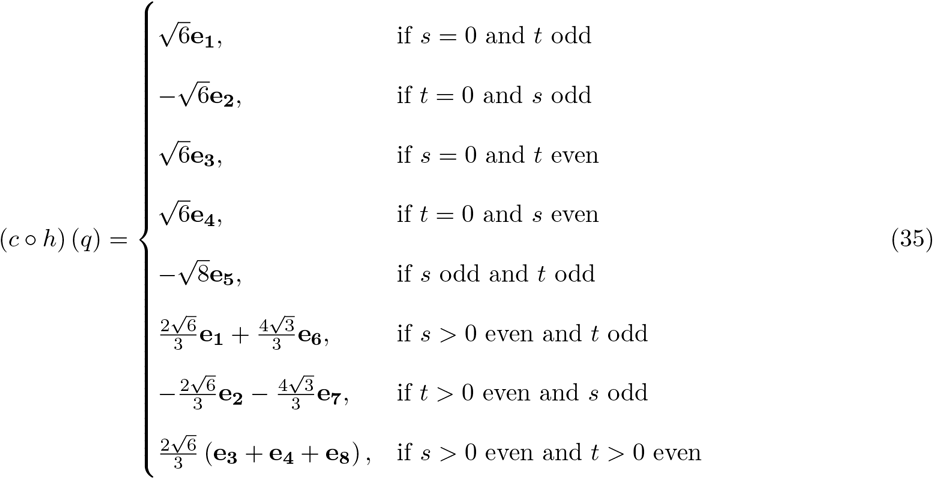

Note that the first inner sum in Equation 34 is a linear combination of basis vectors encoding purely quadratic gradients (**e**_**3**_, **e**_**4**_, **e**_**5**_ and **e**_**8**_), weighted by even trigonometric functions of *θ*. The prefactors on this inner sum are functions that are even in *ϕ*. This inner sum and its prefactor therefore jointly satisfy the similarity relation (*θ, ϕ*) ∼ (*θ* + *π,* 2*π* − *ϕ*) by independently satisfying (*θ*) ∼ (*θ* + *π*) and (*ϕ*) ∼ (2*π* − *ϕ*). Meanwhile, the second inner sum in Equation 34 is a linear combination of basis vectors containing linear gradients (**e**_**1**_, **e**_**2**_, **e**_**6**_ and **e**_**7**_), weighted by odd trigonometric functions of *θ*. The prefactors on this inner sum are functions that are odd in *ϕ*. This inner sum and its prefactor therefore jointly satisfy the similarity relation (*θ, ϕ*) ∼ (*θ* + *π,* 2*π* − *ϕ*), by independently satisfying (*θ*) ∼ −(*θ* + *π*) and (*ϕ*) ∼ −(2*π* − *ϕ*). Since the trigonometric functions of *θ* are coupled to (*x, y*), *θ* controls the rotation of stripes in the image patches, just as in *k*^0^. Similarly, since the prefactors on the inner sums are functions of *ϕ*, *ϕ* controls the relative contribution of quadratic gradients (**e**_**3**_, **e**_**4**_, **e**_**5**_ and **e**_**8**_ in the first inner sum) and linear gradients (**e**_**1**_, **e**_**2**_, **e**_**6**_ and **e**_**7**_ in the second inner sum). Lastly, the boundary conditions for *θ* and *ϕ* in this parameterization of *κ*_*θ,ϕ*_, yield patches with vertical (horizontal) stripes when 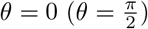, and linear (quadratic) gradients when 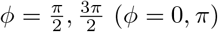 just as in *k*^0^.

A Klein bottle embedding belonging to this parametric family, 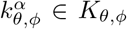, can therefore be specified in terms of a vector *F* = [*N*_*C*_, *N*_*θ*_, *N*_*ϕ*_, *M*_*θ*_, *M*_*ϕ*_] defining its functional form, and a corresponding coefficient vector *α* = [*μ*_*i*_,…, *β*_*i*_,…, *γ*_*i*_,…, *ζ*_*i*_,…, *η*_*i*_,…, *ϑ*_*i*_]. In this parametric family of Klein bottle embeddings, *k*^0^ corresponds to *F* = [0, 0, 0, 2, 1] with *α* = [*η*_1,2,0_, *η*_1,1,1_, *η*_1,0,2_, *ϑ*_1,0,1_, *ϑ*_1,1,0_] = [1, 2, 1, 1, 1]. Note that since curvatures are only computed on the embedding after normalization, *α* is only meaningfully defined up to a multiplicative constant.

#### 4.5.4 Associating Image Patches to a Klein Bottle Embedding

For a given Klein bottle embedding, 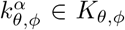, we associated each datapoint *v*_*i*_ (already vectorized and normalized by *u* ◦ *c* ◦ *h*) to the closest point on 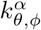 by minimizing the Euclidean distance in ℝ^8^:

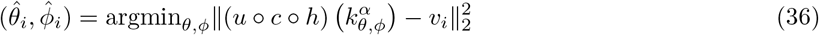

We solved this minimization using the *lsqnonlin* function (‘StepTolerance’=1e-3, ‘FunctionTolerance’=1e-6) in MATLAB, supplying initial conditions corresponding to analytical values for a point on *k*^0^:

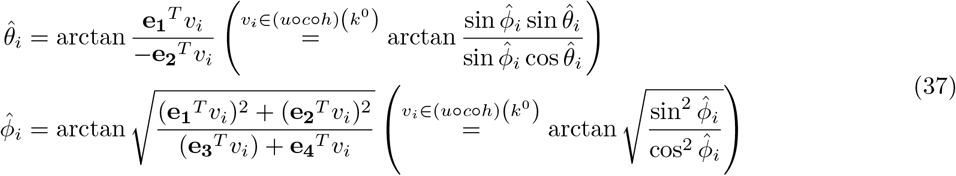

We constrained solutions to 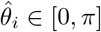 and 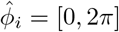 according to the (*θ, ϕ*) similarity relation.

#### 4.5.5 Optimal Klein Bottle Embedding

Let 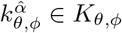 be the Klein bottle embedding that minimizes the sum of Euclidean distances in ℝ^8^ between each image patch and the closest point on the embedding. To determine 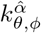 given a functional form *F*, we initialized the coefficient vector 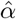 to have zero entries everywhere except for the values used in *k*^0^. We then iterated between optimizing for 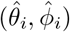 according to Equation 36 and for 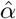 as shown below using least-squares, until convergence:

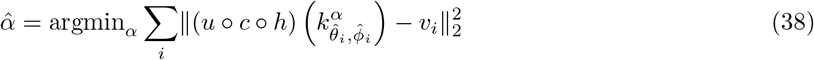

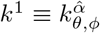 is the optimized Klein bottle embedding corresponding to *F* = [1, 10, 10, 20, 10], for which results are shown in Figures 3H and S3D.

#### 4.5.6 Noisy Klein Bottle Embeddings

The set of *N* ≈ 4.2×10^5^ image patches was associated to *k*^0^ according to the procedure described in Methods Section 4.5.4, yielding 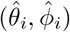 values. Isotropic Gaussian noise of magnitude *sσ* was added element-wise in ℝ^9^ (prior to normalization by *u* ◦ *c*) to 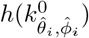, where 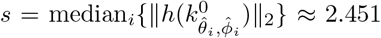. Figures 3F,G and S3A correspond to noise with *σ* = 0.007, 0.01 and 0.03.

#### 4.5.7 Parameters for Curvature Estimation

For all scalar curvature computations on image patch datasets and Klein bottle embeddings, we set *d* = 2 and *N*_*calib*_ = 10^4^. Unless the neighborhoods were manually specified, we used *σ*_*h*_ = 0.1, which yielded a flat distribution of GOF p-values (2.5% of points reported GOF p-values ≤ *α* = 0.05) for the set of *N* ≈ 4.2 × 10^5^ points on *k*^0^ closest to the image patches (shown in Figure 3E).

### 4.6 Details of scRNASeq Datasets

The PBMC dataset provided by 10x Genomics is comprised of *N* = 10194 PBMCs collected from a healthy donor [42]. The mouse gastrulation dataset consists of *N* = 116312 cells collected at nine 6-hour intervals from embryonic day 6.5 to 8.5 [43]. The mouse brain dataset is a benchmark from 10x Genomics consisting of *N* = 1306127 cells collected from the cortex, hippocampus and ventricular zone of two embryonic mice sacked at embryonic day 18 [44].

#### 4.6.1 Preprocessing

For the PBMC dataset, we applied standard preprocessing steps using Seurat v3.1.2 [53] with default function arguments, to extract PC projections and UMAP coordinates ourselves. Specifically, we removed cells where the percentage of transcripts corresponding to mitochondrial genes exceeded 15%, or which had fewer than 500 transcripts. This reduced the number of cells from 10194 to 9385. On this filtered set, we normalized the data (NormalizeData(normalization.method=‘LogNormalize’, scale.factor=10000)), retained the 2000 most variable genes (FindVariableFeatures(selection.method=‘vst’, nfeatures=2000)), and scaled the data (ScaleData). Next, we performed linear dimensionality reduction using PCA down to 50 dimensions (RunPCA(npcs=50)) and generated UMAP coordinates for visualization (RunUMAP(dims = 1:30)). For the gastrulation (brain) dataset, we did not preprocess the data ourselves but instead directly used the 50 (20) PC projections and UMAP (t-SNE) visualization coordinates provided with the dataset. Please refer to [43, 44] for additional details.

#### 4.6.2 Cell Type Annotations

For the PBMC dataset, the AddModuleScore(ctrl=5) function was used to compute the per-cell average expression of marker genes corresponding to seven different cell types [54]. To prepare Figure S4A, each cell was assigned the cell type for which its average marker gene expression was the highest. Cell type annotations for the gastrulation dataset (see Figure S5A) were sourced from Figure 1C of [43]. Cell type annotations for the brain dataset (see Figure S6A) are predicted labels sourced from [55].

#### 4.6.3 Statistical Tests

Here we describe the statistical tests applied to scalar curvatures computed for the scRNAseq datasets.

##### 4.6.3.1 Spatial Precision of Errorbars

Let *m* be the fraction of datapoints with 95% CIs containing the scalar curvatures reported by their respective kNNs. To check whether *m* was significantly larger than chance, we used a permutation test. We randomly assigned the kNN of each datapoint to be one of the *N* datapoints in the dataset and computed *m*. We repeated the procedure *T* = 1000 times to generate an empirical distribution of *m* for the null model of random neighbors. The reported p-value for each *k* is the fraction of the *T* trials for which *m* was greater than the value computed for data. See Figures S4D, S5D and S6D.

##### 4.6.3.2 Sensitivity to Cell Downsampling

To check the sensitivity of the computed scalar curvatures to the average density of cells, we discarded *f* % of cells at random from the ambient space computed using the original set of *N* datapoints, and recomputed scalar curvatures using the same ambient dimension, manifold dimension and neighborhood sizes as for the original dataset (see Methods Section 4.6.4). Let *m* be the fraction of downsampled datapoints with 95% CIs containing the scalar curvatures originally reported. Since the CIs grow as *f* increases, we checked whether *m* was significantly larger than chance by using a permutation test. We randomly paired each of the 95% CIs computed after downsampling, to one of the scalar curvatures reported by the downsampled points for the original dataset, and computed *m*. We repeated the procedure *T* = 1000 times to generate an empirical distribution of *m* for the null model. The reported p-value for each *f* is the fraction of the *T* trials for which *m* was greater than the value computed for data. See Figures S4I, S5I and S6I.

##### 4.6.3.3 Sensitivity to Transcript Downsampling

To check the sensitivity of the computed scalar curvatures to the capture efficiency and sequencing depth of the data, we discarded *f* % of transcripts at random from the single-cell count matrix for the PBMC dataset, then performed the same preprocessing steps described in Methods Section 4.6.1. We recomputed scalar curvatures using the same ambient dimension, manifold dimension and neighborhood sizes as for the original dataset (see Methods Section 4.6.4). Let *m* be the fraction of datapoints with 95% CIs containing the scalar curvatures originally reported. To check whether *m* was significantly larger than chance, we used a permutation test. We randomly paired each of the 95% CIs computed after downsampling transcripts, to one of the scalar curvatures computed for the original dataset, and computed *m*. We repeated the procedure *T* = 1000 times to generate an empirical distribution of *m* for the null model. The reported p-value for each *f* is the fraction of the *T* trials for which *m* was greater than the value computed for data. See Figure S4J.

#### 4.6.4 Parameters for Curvature Estimation

Let the variance explained by the *i*^th^ PC be given by 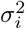 and the cumulative fractional variance of the first *m* PCs by 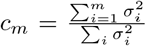. For each dataset, we selected the ambient dimension as *n* = argmax*m*{*c*_*m*_|*c*_*m*_ ≤ 0.8}, the manifold dimension as *d* = argmax*m*{*c*_*m*_|*c*_*m*_ ≤ 0.64}, and considered the global length scale to be *L* = 3*σ*_*d*_. (*n, d, L*) = (8, 3, 18.3), (11, 3, 19.1) and (9, 5, 24.9) for the PBMC, gastrulation and brain datasets respectively. For the three datasets, we computed scalar curvatures for manifold dimensions *d* − 1, *d* and *d* + 1. It was not always possible to select *σ*_*h*_ for each dataset and manifold dimension, so that the distribution of GOF p-values was flat, according to our usual heuristic. For consistency, we therefore picked *σ*_*h*_ so that 1/3 of points had GOF p-values ≤ *α* = 0.05. For manifold dimension (*d* − 1*, d, d* + 1), *σ*_*h*_ = (0.031, 0.041, 0.045), (0.036, 0.044, 0.053) and (0.034, 0.050, 0.055) for the PBMC, gastrulation and brain datasets respectively.

## 5 Acknowledgements

DS was funded in part by the Natural Sciences and Engineering Research Council of Canada (NSERC PGSD2-517131-2018). SW was supported by NCI U54-CA225088 and NIH NIGMS T32 GM008313. DS and SH acknowledge funding from NIH NIGMS R00GM118910, U19 Systems Immunology Pilot Project Grant at Harvard University, and the Harvard University William F. Milton Fund. The authors would like to thank Peter Kharchenko and Allon Klein for helpful discussions. Portions of this research were conducted on the O2 High Performance Compute Cluster, supported by the Research Computing Group, at Harvard Medical School. See http://rc.hms.harvard.edu for more information.

## 6 Data and Code Availability

The *van Hateren* IML dataset is available at http://bethgelab.org/datasets/vanhateren and was loaded according to the instructions there. The PBMC dataset is available at https://support.10xgenomics.com/single-cell-gene-expression/datasets/4.0.0/Parent_NGSC3_DI_PBMC. The gastrulation dataset can be retrieved using instructions found at https://github.com/MarioniLab/EmbryoTimecourse2018. The brain dataset is available at https://support.10xgenomics.com/single-cell-gene-expression/datasets/1.3.0/1M_neurons. The software package described here to compute scalar curvature is available at https://gitlab.com/hormozlab/ManifoldCurvature. All code and instructions to reproduce the numerics and figures in this study will be made available upon publication.

## 7 Supplementary Figures

**Figure S1:**
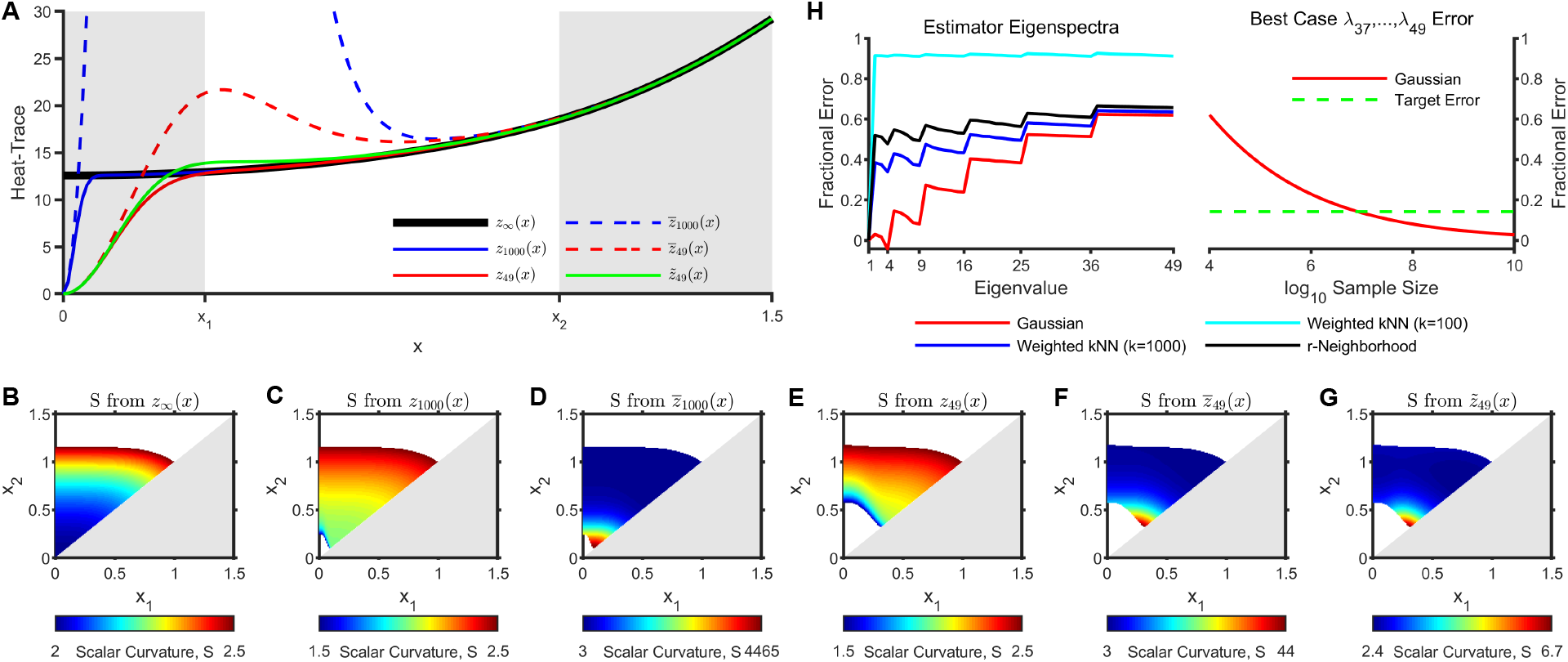
The scalar curvature of 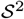 is poorly estimated using the Laplace-Beltrami operator. **(A)** The heat-trace with *m* terms, (*zm*(*x*) in Equation 3) is shown for *m* = ∞ (black), *m* = 1000 (solid blue) and *m* = 49 (solid red), when evaluated with analytical eigenvalues for 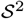. Empirical eigenvalues were obtained by uniformly sampling *N* = 10^4^ points from 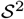 (see Figure 1A; Methods Section 4.4.1.1) and estimating the Laplace-Beltrami (LB) operator using Equations 13-15. The heat-trace evaluated using these empirical eigenvalues, 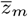, is shown for *m* = 1000 (dashed blue) and *m* = 49 (dashed red). The heat-trace evaluated using eigenvalues obtained by interpolating between the analytical and empirical values 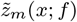 in Equation 11) is shown for *m* = 49 and *f* = 0.23 (solid green). *f* signifies that the fractional error of the interpolated eigenvalues is reduced by 1 − *f* relative to the empirical eigenvalues. *f* = 0 corresponds to the analytical eigenvalues while *f* = 1 corresponds to the empirical eigenvalues. The white region bounded by [*x*_1_*, x*_2_] indicates a candidate interval over which to fit a heat-trace to a quadratic in order to extract an estimate for the scalar curvature (see Equations 2-4; Methods Section 4.2.1). On the one hand, since the knee of *zm*(*x*) shifts to the left as *m* increases (i.e. *z*_*m*_(*x*) converges from ∞), larger *m* results in more intervals for which *zm*(*x*) well-approximates *z*_*∞*_(*x*) and will therefore yield accurate scalar curvature estimates. On the other hand, 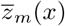 becomes a worse estimator for *zm*(*x*) as *m* increases. **(B)** Scalar curvatures estimated by fitting *z*_*∞*_(*x*) to a quadratic over different intervals [*x*_1_*, x*_2_] as defined in (A). Scalar curvatures are shown in color for intervals yielding accurate estimates (*S* ∈ [1.5, 2.5]). This colored region corresponds to *D*_*∞*_. **(C)** As in (B) but with estimates obtained by fitting a quadratic to *z*_1000_(*x*). The colored region corresponds to *D*_1000_. By inspection, *D*_1000_ ⊂ *D*_*∞*_. **(D)** Scalar curvatures estimated by fitting 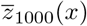 to a quadratic over each interval in *D*_1000_. Though *D*_1000_ was constructed using only intervals which yielded an accurate scalar curvature estimate when analytical eigenvalues were used in the heat-trace, no interval in *D*_1000_ yields an accurate scalar curvature estimate when the same number of empirical eigenvalues are used in the heat-trace instead. **(E)** As in (B) but with estimates obtained by fitting a quadratic to *z*_49_(*x*). The colored region corresponds to *D*_49_. By inspection, *D*_49_ ⊂ *D*_1000_ **(F)** As in (D) but with estimates obtained by fitting 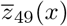 to a quadratic over each interval in *D*_49_. No estimate is accurate just as in (D). **(G)** As in (F) but with estimates obtained by fitting 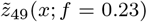 to a quadratic over each interval in *D*_49_. *f* = 0.23 was chosen so that half the intervals in *D*_49_ yield an accurate scalar curvature estimate. **(H)** (Left) The fractional error in the first 49 empirical eigenvalues of the LB estimator from (A) is shown in red. This operator was computed using the Gaussian kernel (*Wg* in Equation 15). Eigenvalues 37-49 have a fractional error of 60%. The fractional error of the eigenvalues of LB estimators computed on the same *N* = 10^4^ points but using the weighted kNN and *r*-neighborhood kernels (*W*_*kNN*_ and *W*_*r*_ respectively in Equation 16) is also plotted. Positive error indicates under-estimation. (Right) Projected fractional error for eigenvalues 37-49 of the LB estimator with Gaussian kernel computed using a larger sample size (*N*). The projection is based on the convergence rate given in Theorem 1 of [30], assuming that the big-*O* bound is sharp at *N* = 10^4^ for eigenvalues 37-49. The dashed green line corresponds to the 14% fractional error needed for scalar curvatures to be accurately estimated for half the intervals in *D*_49_. This corresponds to *f* = 0.23 in (G) since 60% × *f* = 14%. For the LB estimator computed using the Gaussian kernel, achieving this fractional error requires *N*10^7^. Since LB estimators computed using the other kernels have the same convergence rate but larger fractional error at *N* = 10^4^, these estimators would require even larger *N* to achieve the desired 14% fractional error.

**Figure S2:**
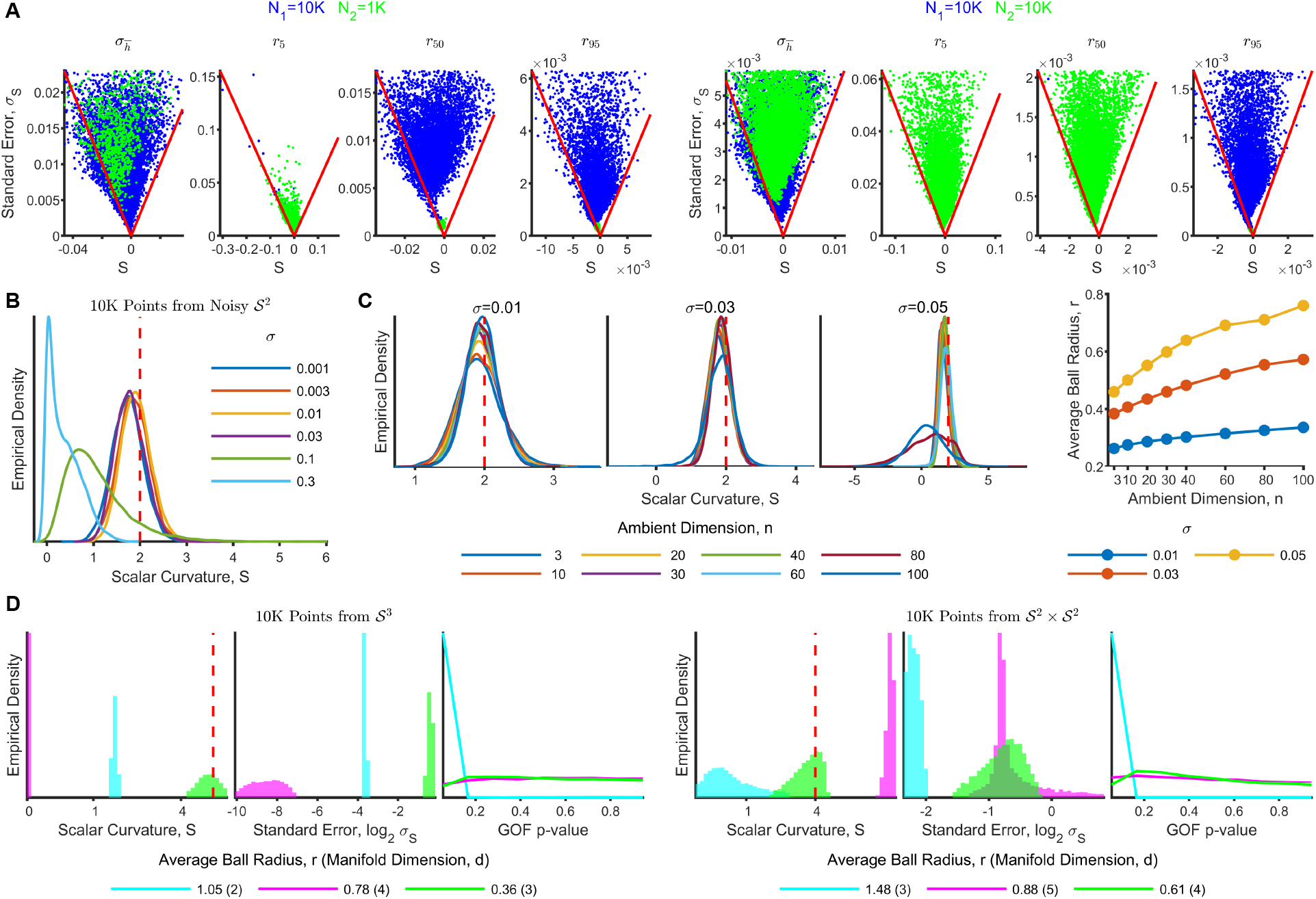
Sensitivity of algorithm to real-world confounders. **(A)** (Left) A dataset with a sparse periphery and a dense core was formed by uniformly sampling *N*_1_ = 10^4^ points from the 3-dimensional cube of side-length 10, 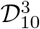, and *N*_2_ = 10^3^ points from the 3-dimensional cube of side-length 1, 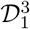 (see Methods Section 4.4.1.4). These points were embedded in ℝ^11^ and padded with isotropic Gaussian noise of magnitude *σ* = 0.01 in the normal directions. Scalar curvatures (*S*) were computed on this dataset of *N*_1_ + *N*_2_ points by setting *σ*_*h*_ and are plotted against their standard errors (*σ*_*S*_) in the leftmost panel. Curvature computations were also performed at fixed length scales corresponding to the 5, 50 and 95%-ile values for neighborhood size (left to right) used in the leftmost panel (*r* = 0.54, 0.90 and 1.22 respectively). Here, points for which the chosen *r* led to neighborhoods with insufficient points for regression are not shown. For large length scales, all points in the dense region are able to report curvatures but are crowded into the apex of the plots. The *N*_1_ (*N*_2_) sparse (dense) points are shown in blue (green). Points enclosed by the red lines have 95% CIs including the true value of zero. The right four panels show analogous results when *N*_2_ = 10^4^. Here the the 5, 50 and 95%-ile values for neighborhood size are *r* = 0.37, 0.63 and 1.42 respectively. See Methods Section 4.4.2.1. **(B)** Distribution of scalar curvatures computed for *N* = 10^4^ points uniformly sampled from ^2^ ℝ^3^ and convoluted with isotropic Gaussian noise of magnitude 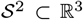. Noise confounds accurate scalar curvature computation when *σ* is roughly 10% of the sphere’s radius. The deviation of the estimated scalar curvatures from the true value of 2 (shown as a dashed red line) for *σ* ≥ 0.1 reflects the nontrivial geometry of a manifold convoluted by noise. See Methods Section 4.4.2.2. **(C)** (Left) *N* = 10^4^ points were uniformly sampled from 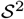 and embedded in ℝ^*n*^. Isotropic Gaussian noise of magnitude *σ* was applied to each of the *n* ambient dimensions. Scalar curvatures computed by keeping *σ*_*h*_ fixed for all *n* and *σ*, recapitulated the true value of 2 (shown as dashed red lines) for *n* ≤ 80 and *σ* ≤ 0.05. (Right) The neighborhood size (*r*) necessary to attain *σ*_*h*_ is less sensitive to changes in *n* than changes in *σ*. See Methods Section 4.4.2.3. **(D)** *N* = 10^4^ points were uniformly sampled from (left) 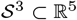 convoluted with isotropic Gaussian noise in the ambient space with *σ* = 0.01 and (right) 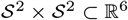. To investigate the effects of choosing the manifold dimension, *d*, differently than the true value, *d*\*, *σ*_*h*_ was kept fixed, and scalar curvatures were computed for *d* = *d*\* − 1 (cyan), *d* = *d*\* + 1 (magenta) and *d* = *d*\* (green). The panels show the distribution of (left to right) scalar curvatures (*S*), standard errors (*σ*_*S*_) and GOF p-values. The true value of the scalar curvature (at *d* = *d*\*) is constant across both manifolds and shown as a dashed red line. The average neighborhood size (*r* averaged over all points) is much larger for both *d* = *d*\* − 1 and *d* = *d*\* + 1 than for *d* = *d*\* as shown in the legend. For the same *σ*_*h*_, *d* = *d\** 1 also leads to a more skewed distribution of GOF p-values relative to *d* = *d*\*, while the distribution for *d* = *d*\* + 1 is still flat. See Methods Section 4.4.2.4.

**Figure S3:**
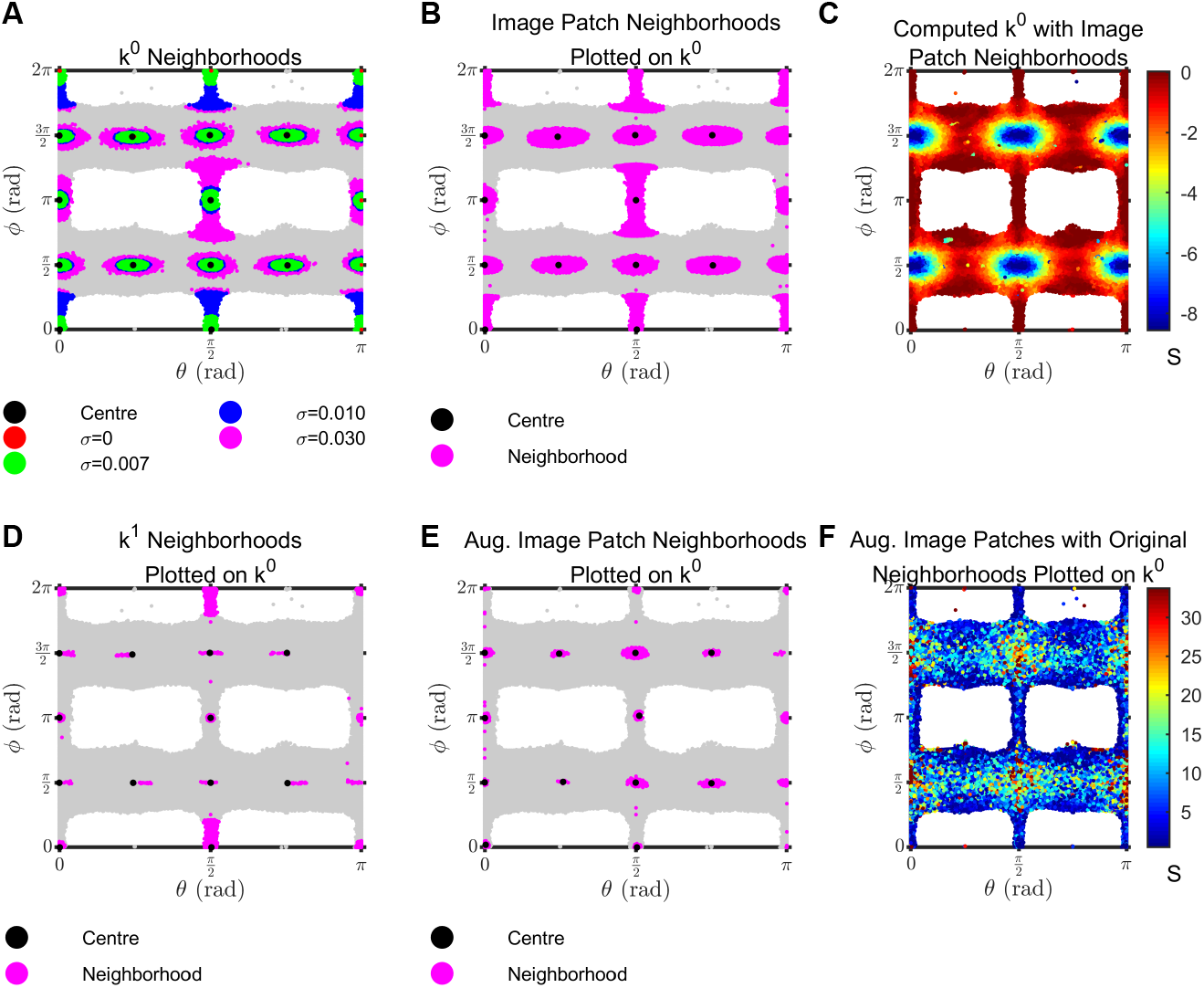
Additional details of the image patch dataset and Klein bottle embeddings (related to Figure 3). **(A)** To compute scalar curvatures for Figure 3E, each image patch was associated to the (*θ*_0_*, ϕ*_0_) coordinates of the closest point on *k*^0^. Here we select a handful of these associated points on *k*^0^ (shown in black) and visualize how neighborhoods chosen in ℝ^8^ to compute scalar curvatures for Figure 3E appear in (*θ*_0_*, ϕ*_0_) coordinates (shown in red). When noise of increasing magnitude, *σ*, is added to the set of closest points on *k*^0^ (see Methods Section 4.5.6), the neighborhood size at each point grows until *σ_h_* is attained. **(B)** As in (A), but showing neighborhoods used in computing the scalar curvatures in Figure 3D for the image patch dataset. Note the close correspondence in neighborhood size with *σ* = 0.03 in (A). **(C)** Scalar curvatures computed for the set of closest points (*θ*_0_*, ϕ*_0_) on *k*^0^ as in Figure 3E, but using the same neighborhood sizes determined for the image patch dataset shown in Figure 3D, some of which are visualized in (B). **(D)** As in (A) but showing neighborhoods used in computing the scalar curvatures in Figure 3H for the set of closest points on *k*^1^. Neighborhoods are visualized on (*θ*_0_*, ϕ*_0_) coordinates instead of (*θ*_1_*, ϕ*_1_) coordinates for ease of comparison. **(E)** As in (B) but showing neighborhoods used in computing the scalar curvatures in Figure 3J for the augmented image patch dataset. **(F)** Scalar curvatures computed for the augmented image patch dataset with *N* 1.3 10^8^ points as in Figure 3J, but using the same neighborhood sizes determined for the original image patch dataset with *N* 4.2 10^5^ shown in Figure 3D and (B). Note the close correspondence with Figure 3D.

**Figure S4:**
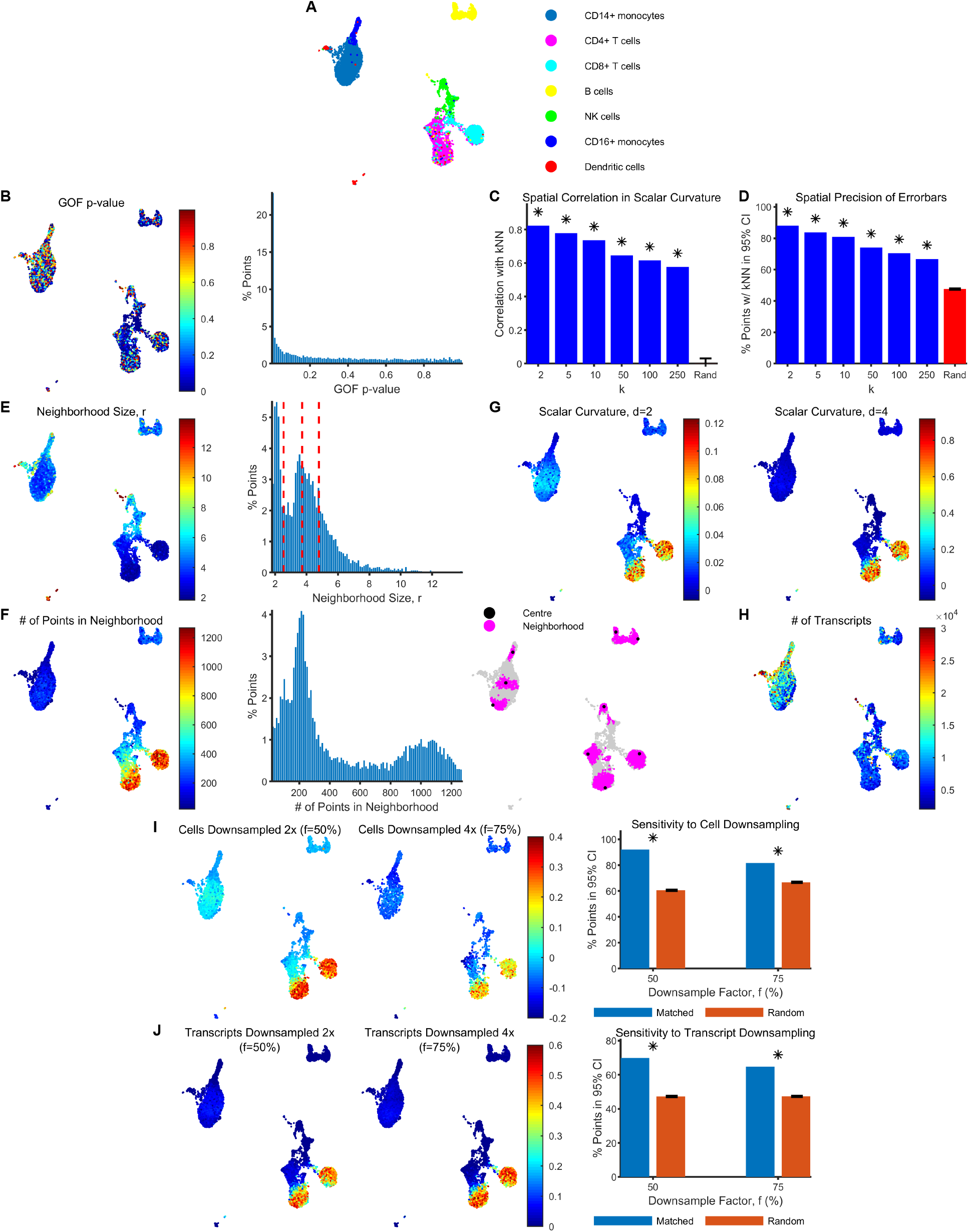
Additional details of the PBMC scRNAseq dataset (related to Figure 4). **(A)** Cell types overlaid onto UMAP coordinates and sorted in decreasing order of abundance in the legend. Cells were annotated as described in Methods Section 4.6.2. **(B)** A goodness-of-fit p-value was computed for each point by applying Mardia’s test to the residuals obtained from fitting the neighborhood around the point to a quadratic function (see Methods Section 4.3.3). These p-values are visualized on UMAP coordinates corresponding to each point (left) and their empirical distribution is shown using a histogram (right). Small p-values suggest that the residuals are non-normal so that approximating local neighborhoods as quadratic may not be valid. **(C)** Pearson correlation between the scalar curvature reported by each point and its *k*^th^-nearest neighbor (kNN) for different *k* (shown in blue). The red bar shows the mean and standard deviation of the Pearson correlation when neighbors are chosen randomly over 1000 trials (\**p* < 10^*−*6^). **(D)** The percentage of points with 95% CIs containing the scalar curvatures reported by their respective kNNs (shown in blue). The red bar shows the mean and standard deviation of this percentage when neighbors are chosen randomly over 1000 trials (\**p* < 0.001; see Methods Section 4.6.3.1). **(E)** The neighborhood size (*r*) used for computing scalar curvature at each point, overlaid onto UMAP coordinates (left) and a corresponding histogram of the empirical distribution (right). The dashed red lines correspond to the 25, 50, and 75%-ile values of *r*(*p*) used for computing scalar curvatures at fixed neighborhood sizes for Figure 4C. See Methods Section 4.3.2. **(F)** The number of points in each neighborhood (corresponding to the neighborhood sizes in (E)) overlaid onto UMAP coordinates (left) and a corresponding histogram of the empirical distribution (middle). (Right) The set of neighbors used for computing scalar curvature (purple) is visualized on UMAP coordinates for a handful of points (black). **(G)** Scalar curvatures were computed for manifold dimension *d* − 1 (left) and *d* + 1 (right). They are plotted here on UMAP coordinates after smoothing over the same set of *k* = 250 neighbors used in Figure 4A. See Methods Section 4.6.4. **(H)** The total number of transcripts observed in each cell overlaid onto UMAP coordinates. **(I)** Scalar curvatures were computed after downsampling the number of cells in the ambient space by a factor of 2 (left) and 4 (middle), using the same ambient dimension, manifold dimension and neighborhood sizes determined for the original dataset. They are plotted here on UMAP coordinates after smoothing over the same set of neighbors (which survive downsampling) used in Figure 4A. (Right) The percentage of points in the downsampled datasets with a 95% CI containing the originally reported scalar curvature (blue), and likewise for a negative control obtained by randomly pairing 95% CIs and originally reported scalar curvatures for points in the downsampled dataset (red). Errorbars for the negative control are the standard deviation of this percentage over 1000 trials with different random pairings (\**p* < 0.001; see Methods Section 4.6.3.2). Scalar curvatures were computed after downsampling the number of transcripts by a factor of 2 (left) and 4 (middle), using the same ambient dimension, manifold dimension and neighborhood sizes determined for the original dataset. They are plotted here on UMAP coordinates after smoothing over the same set of *k* = 250 neighbors used in Figure 4A. (Right) The percentage of points in the downsampled datasets with a 95% CI containing the originally reported scalar curvature (blue), and likewise for a negative control obtained by randomly pairing 95% CIs and originally reported scalar curvatures for points in the downsampled dataset (red). Errorbars for the negative control are the standard deviation of this percentage over 1000 trials with different random pairings (\**p* < 0.001; see Methods Section 4.6.3.3).

**Figure S5:**
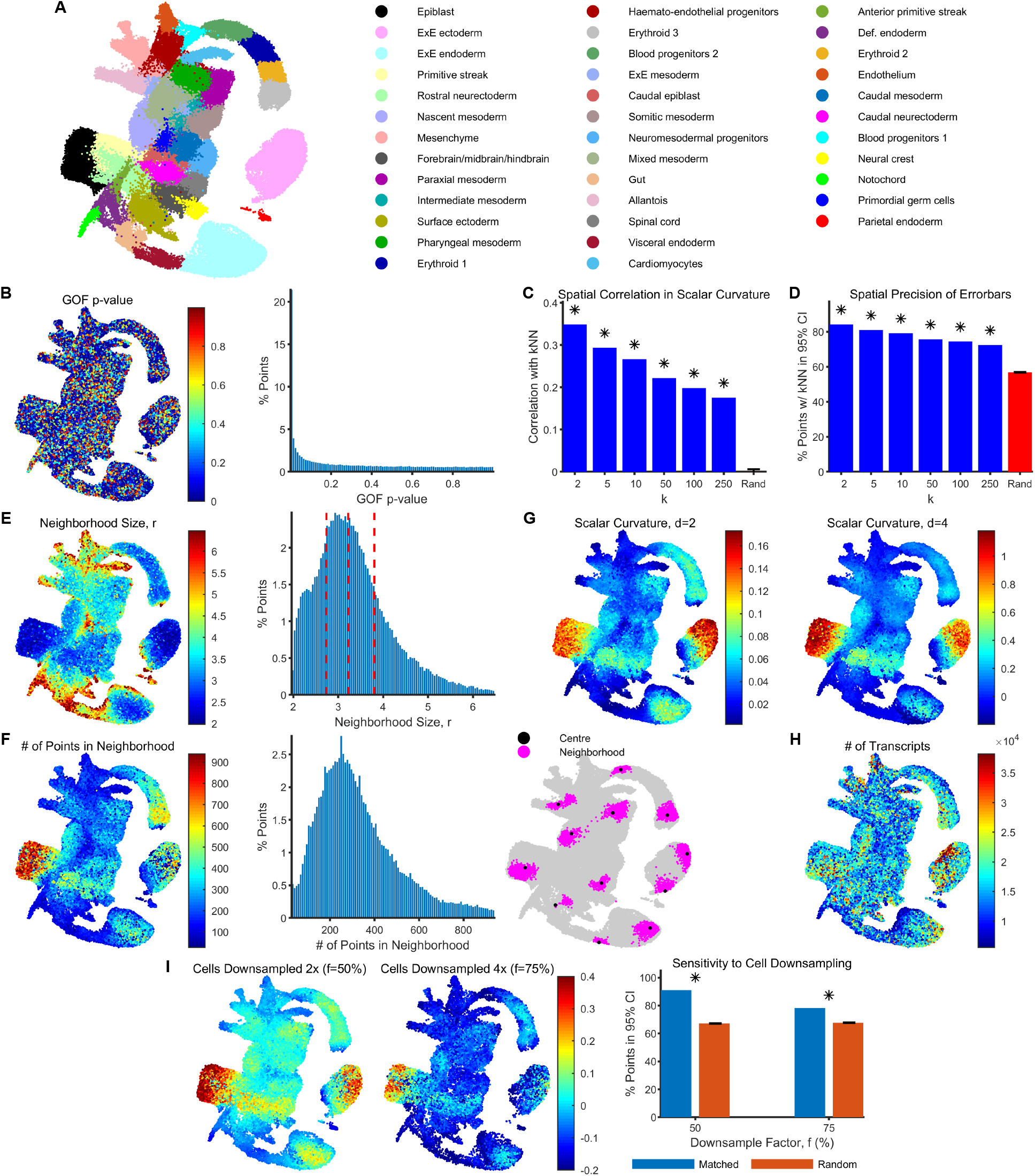
Additional details of the gastrulation scRNAseq dataset (related to Figure 4). Panels as in Figure S4.

**Figure S6:**
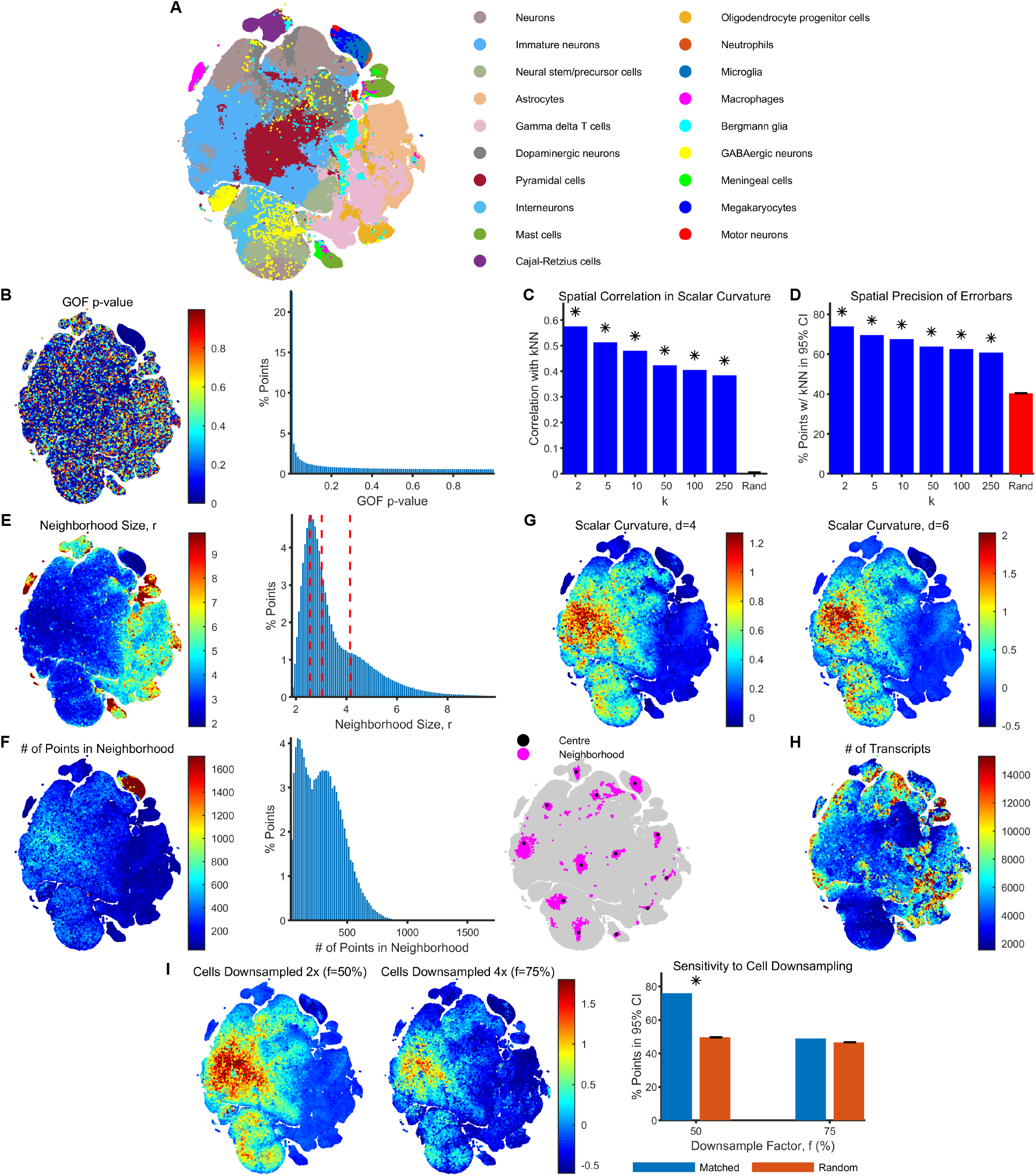
Additional details of the brain scRNAseq dataset (related to Figure 4). Panels as in Figure S4 but with t-SNE instead of UMAP plots.

